# Emergent Elasticity in the Neural Code for Space

**DOI:** 10.1101/326793

**Authors:** Samuel Ocko, Kiah Hardcastle, Lisa Giocomob, Surya Ganguli

## Abstract

Upon encountering a novel environment, an animal must construct a consistent environmental map, as well as an internal estimate of its position within that map, by combining information from two distinct sources: self-motion cues and sensory landmark cues. How do known aspects of neural circuit dynamics and synaptic plasticity conspire to accomplish this feat? Here we show analytically how a neural attractor model that combines path integration of self-motion cues with Hebbian plasticity in synaptic weights from landmark cells can self-organize a consistent map of space as the animal explores an environment. Intriguingly, the emergence of this map can be understood as an elastic relaxation process between landmark cells mediated by the attractor network. Moreover, our model makes several experimentally testable predictions, including: (1) systematic path-dependent shifts in the firing field of grid cells towards the most recently encountered landmark, even in a fully learned environment, (2) systematic deformations in the firing fields of grid cells in irregular environments, akin to elastic deformations of solids forced into irregular containers, and (3) the creation of topological defects in grid cell firing patterns through specific environmental manipulations. Taken together, our results conceptually link known aspects of neurons and synapses to an emergent solution of a fundamental computational problem in navigation, while providing a unified account of disparate experimental observations.

---

How might neural circuits learn to create a long term map of a novel environment and use this map to infer where one is within the environment? This pair of problems are challenging because of their nested, chicken and egg nature. To localize where one is in an environment, one first needs a map of the environment. However, in a novel environment, no such map is yet available, so localization is not possible. Instead, neural circuits must create a map over time, through exploration in a novel environment, without initially having access to any global estimate of position within the environment. This chicken and egg problem is known in the robotics literature as Simultaneous Localization and Mapping (SLAM) (1).

Here we explore how known aspects of neural circuit dynamics and synaptic plasticity can conspire to self-organize, through exploration, a neural circuit solution to the problem of creating a global, consistent map of a novel environment. In particular, neural circuits receive two fundamentally distinct sources of information about position: (1) signals indicating the speed and direction of the animal, which can be path-integrated over time to update the animal’s internal estimate of position, and (2) sensory cues from salient, fixed landmarks in the environment. To create a map of the environment, neural circuits must combine these two distinct information sources in a self-consistent fashion so that sensory cues and selfmotion cues are always in co-register.

For example, consider the act of walking from landmark A to landmark B. Sensory perception of landmark A triggers a pattern of neural activity, and subsequent walking from A to B evolves this activity pattern, through path integration, to a final pattern. Conversely, sensory perception of landmark B itself triggers a neural activity pattern. Any circuit that maps space must obey a fundamental self-consistency condition: the neural activity pattern generated by perception of A, followed by path integration from A to B, must match the neural activity pattern triggered by perception of B alone. Only in this manner can neural activity patterns be in one to one correspondence with physical positions in space, and become independent of the past trajectory used to reach any physical location.

In the following, we develop an analytic theory for how neuronal dynamics and synaptic plasticity can conspire to self-organize such a self-consistent neural map of space upon exploration through a novel environment. Moreover, our analytic theory makes experimentally testable predictions about neural correlates of space. Indeed, many decades of recordings in multiple brain regions have revealed diverse neural correlates of spatial maps in the brain. In particular, the medial entorhinal cortex (MEC) contains neurons encoding for direction, velocity, landmarks, as well as grid cells exhibiting striking firing patterns reflecting an animal’s spatial location (2–6). Moreover, the geometry of these firing patterns depends on the shape of the environment being explored (7–10). In particular, these grid firing patterns can be deformed in irregular environments (11, 12), in a manner evocative of deformations of solids forced into into an irregular container, suggesting a mechanical model for these deformations (13–15). Also, these firing patterns are not simply driven by current sensory cues; there is evidence for path integration (16–18) in that firing patterns appear almost immediately (2), phase differences are preserved across environments (19), firing patterns become noisier the longer an animal has spent away from a landmark (20, 21), and can be shifted depending on which landmark the animal has most recently encountered (22, 23).

Despite this wealth of experimental observations, no mechanistic circuit model currently explains how known aspects of neuronal dynamics and synaptic plasticity can conspire to learn, through exploration, a self-consistent internal map of a novel environment that both behaves like a deformable medium, and also retains, at higher-order, some knowledge of recently encountered landmarks. Here, we show how an attractor network that combines path integration of velocity with Hebbian learning (22, 24, 25) of synaptic weights from landmark cells, can self-organize to generate all of these outcomes. Intriguingly, a low dimensional reduced model of the combined neuronal and synaptic dynamics provides analytical insight into how self-consistent maps of the environment can arise through an emergent, elastic relaxation process involving the synaptic weights of landmark cells.

## Model reduction of an attractor network coupled to sensorimotor inputs

Our theoretical framework assumes the existence of three interacting neural components: (1) an attractor network capable of realizing a manifold of stable neural activity patterns, (2) a population of velocity-tuned cells that carry information about the animal’s motion, and (3) a population of sensory driven landmark cells that fire if and only if fire animal is in a particular region of space. Our goal will be to understand how the three populations can interact together and self-organize through synaptic plasticity, sculpted by experience, to create a self-consistent internal map ol tine environment. Here, we describe the neuronal and synaptic dynamics of each component in turn, as well as describe a model reduction approach to obtain a low dimensional reduced description of the entire plastic circuit dynamics. Our low dimensional description provides insight into how self-consistency of the neural map emerges naturally through an elastic relaxation process between landmarks.

### A manifold of stable states from attractor netwosk dynamics

We first consider a one-dimensional attrnetor network consisting of a large population of neurons whose connectivity is determined by their position on an abstract ring, as in Fig. 1. For analytical simplicity, we take a neural field approach (26), so that position on the ring of neurons is described by a continuous coordinate *u*, with the firing rate of a neuron at position *u* given by *s*(*u*). Each neuron interacts with neighboring neurons through a translation invariant connectivity, yielding the dynamics

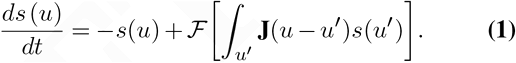

**Fig. 1.**
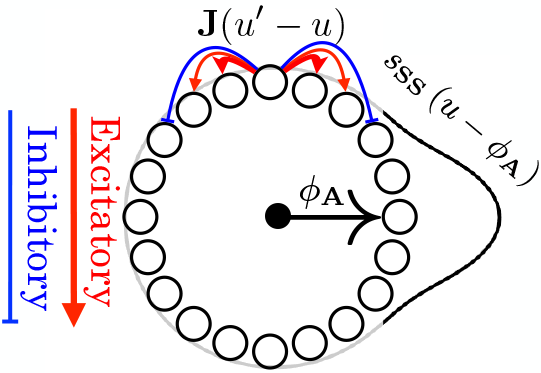
Schematic of a ring attractor with short-range excitation (red arrows) atd longer range inhibition (blue arrow/s). This yieldu a 1S family af bump-attractor stntes *s*_**SS**_ (*u* − *ϕ*_**A**_), which are mapped onto a single periodic variable *ϕ*_**A**_ repre. senting the peak of the bump pattern.

Here **J**(*u* − *u*′) defines the synaptic weight from a cell at position *u′* to a cell at *u*, and 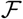 is a nonlinearity. We will refer to these dynamics as *ds/dt* = Dyn[*s*]. Many appropriate choices of **J** and 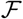, corresponding for example to short range excitation and long range inhibition, will yield a family of stable, or steady state, localized bump activity patterns *s*_**SS**_ (*u* − *ϕ*_**A**_), parameterized by the position of their peak: *ϕ*_**A**_ (27, 28). This one-dimensional family of stable bump activity patterns can itself be thought of as ring of stable firing patterns in the space (if all possible firing patterns. Just as *u* indexes a family of neurons on the neural sheet, the coordinate *ϕ*_**A**_ indexes the different stable neural activity patterns, with a particular value of *ϕ*_**A**_ corresponding to a stable bump on tire neural ring centered at coordinate *u* = *ϕ*_**A**_. For simplicity we oet units such that the coordinate *u* along the neural ring, and the coordinate *ϕ*_**A**_ along the ring of stable attractor patterns are both periodic variables defined modulo 2π. Thus *u* and *ϕ*_**A**_ are phase veriables denoting position along the neural ring ond ring ot bump) attractor patterns respectively.

### Motions along the attractor manifold due to external inputs

So far, the attractor network described above has a ring of stable bump activity patterns parameterized by the periodic coordinate *ϕ*_**A**_, but these neural activity patterns are as yet unanchored to physical space. We will eventually show how to anchor the coordinate *ϕ*_**A**_ along the attractor manifold to the actual position of the animal in physical space. However, in order to appropriately form such an internal map of position, and thereby map the environment, the attractor state must be influenced by external inputs from both velocity and landmark sensitive cells in a self-consistent manner. We first derive a reduced description for how a general external feedforward input to the attractor network modifies its dynamics.

Suppose the attractor network is at one of its steady state bump patterns *s*_**SS**_ (*u* − *ϕ*_**A**_) centered at *u* = *ϕ*_**A**_. Further suppose that each neuron at position *u* on the neural ring experiences an external additive input current *ϵ*Pert (*u* − *ϕ*^**p**^) that is centered, or localized on the neural ring around some other location *u* = *ϕ*^**p**^. The neural dynamics of the attractor in Eq. 1, in response to this additive external input is then modified to:

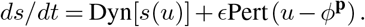

When *ϵ* is small, the external inputs are weak relative to the recurrent inputs that determine the shape of the bump pattern. In this situation, the evolving firing rates will be confined to the 1D manifold of steady states *s*_SS_ (*u* − *ϕ*_**A**_). In essence, a small excitatory perturbation Pert (*u* − *ϕ*^**p**^) centered at position *u* = *ϕ*^**p**^ on the neural ring, will translate the stable bump pattern towards the perturbation, *without* changing its shape. Therefore, to track the entire dynamics of the network, we do not need to track the firing rate of every neuron; we need only track the time dependent position of the peak of the activity bump, *ϕ*_**A**_(*t*). Thus we can reduce the entire high dimensional neural dynamics to a low dimensional effective scalar dynamics:

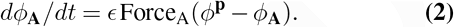

In App. A, we show how to analytically compute the force law Force_**A**_(*ϕ*^**p**^ − *ϕ*_**A**_) governing the velocity of the bump peak, in terms of the shape of the bump *s*_**SS**_ (*u* − *ϕ*_**A**_) and the shape of the additive input perturbation Pert (*u* − *ϕ*^**p**^). However, for the particular external inputs we consider below, the qualitative structure of the force law as a function of the input perturbation will be highly intuitive.

### Path integration through conjunctive position velocity inputs

Following (27, 28), we achieve path integration by coupling the attractor network to conjunctive position and velocity-tuned cells such that east (west) movement-selective cells form feedforward synapses into the attractor network that are shifted in the positive (negative) *u* direction (Fig. 2A, B). We can use our model reduction framework via Eq. 2 to show analytically (App. B) that this choice of connectivity leads to path integration:

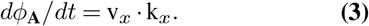

Here, k_*x*_ is a constant of proportionality that relates animal velocity to the rate of phase advance in the attractor network(k_*x*_ = 2π/Field Spacing). Solving Eq. 3 allows us to recover path integration where the resulting integrated attractor phase is *only* a function of current position **r**(*t*):

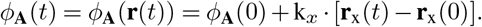

Thus the connectivity of the conjunctive position velocity cells in Fig. 2A, B ensure that as the mouse moves east (west) along a 1D track, the attractor phase moves clockwise (counterclockwise), at a speed proportional to velocity. The collection of neurons in the attractor then trace out periodic firing patterns as a function of spatial position, all with the same period but different phases.

**Fig. 2.**
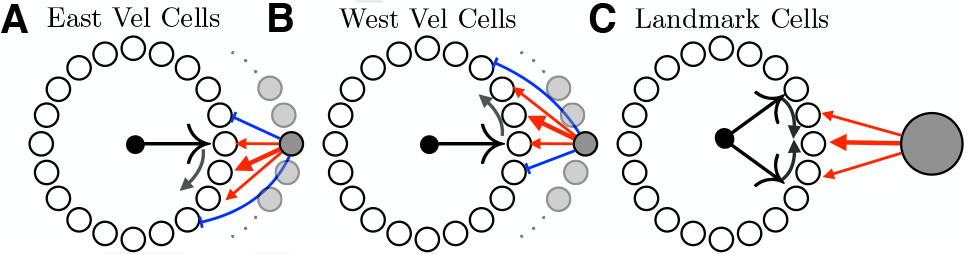
**A)** When the animal moves east, east-conjunctive cells with biased outgoing) connections move the attractor pattern in the positive *u* direction. **B)** When the animal moves west, the attractor pattern is moved in the negative *u* direction. **C)** Schematic of a landmark cell correcting the the attractor bump (Eq. 4). A single landmark cell will pull the peak of the bump pattern towards the peak of its efferent synaptic strength profile.

However, even though these 1D grid cell firing patterns are now a function of physical space, they still are not yet anchored to the environment. There is as yet no mechanism to set the phase of each cell relative to landmarks, and indeed these grid patterns rapidly decohere without anchoring to landmarks, as demonstrated experimentally (20, 29). Coupling the attractor network to landmark-sensitive cells can solve this problem.

### Landmark Cells

We model each landmark cell *i* as purely sensory driven cell with a firing rate that depends on location through Firing_*i*_(*t*) = H_*i*_(**r**(*t*)). Here H_*i*_(**r**) is the firing field of the landmark cell. An example of a landmark cell could, for example, be an entorhinal border cell (4). Every landmark cell forms feed-forward connections onto each cell in the attractor network at ring position *u* with a synaptic strength W_*i*_(*u*).

Consider for example a single landmark cell whose synaptic strength W(*u*) as a function of position *u* on the neural ring consists of a single bump centered at a particular location *u* = *ϕ*^L^(Fig. 2C). Through our model reduction framework of Eq. 2, if this landmark cell fires, then it will exert a force on any other attractor bump pattern *s*_**SS**_(*u* − *ϕ*_**A**_) centered at *u* = *ϕ*_**A**_, through:

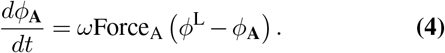

Here we have introduced *ω* as a parameter that controls how strongly landmark cells influence the attractor phase. In essence, when each landmark cell fires, it forces the the attractor state *ϕ*_**A**_ to flow towards the phase *ϕ*^L^ corresponding to the location of its maximal outgoing synaptic strength. An attractor phase *ϕ*_**A**_ that is smaller (larger) than the landmark cell synapses’ peak location *ϕ*^L^ will increase (decrease) and settle down at *ϕ*^L^ (Fig. 2C). Indeed for general synaptic strength patterns peaked at *u* = *ϕ*^L^, the force law will have the same qualitative features as Force_A_ (*ϕ*^L^ − *ϕ*_**A**_) = sin(*ϕ*^L^ − *ϕ*_**A**_) (App. B.1).

However there is, as of yet, no mechanism to enforce consistency between the path integration dynamics of the attractor network in the absence of landmarks in Eq. 3, with the driving force exerted by a landmark cell to a particular phase *ϕ*^L^ through Eq. 4. We next introduce Hebbian plasticity of efferent landmark cell synapses during exploration while *both* path integration and landmark cells are active. We then show how such plasticity yields a precise mechanism for the self-consistency required of any spatial map forming circuit.

### Hebbian Learning Between Landmark Cells and Attractor Networks

We assume that each synapse W_*i*_(*u*) from a landmark cell *i* to an attractor cell at position *u* undergoes Hebbian plasticity with some weight decay, thereby learning to reinforce attractor patterns that are active when the landmark cell fires. Moreover, we assume the dynamics of plasticity varies slowly, over a timescale T that is much longer than the timescale *t* over which explorations occur. Then Hebbian learning drives synaptic strengths towards the long-time average of attractor states that occur during landmark cell firing through

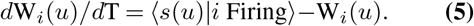

Assuming the effect of landmark cells on the attractor network is strong enough to affect the position of the bump patterns, but not strong enough to change their shape, then the long term average 〈*s*(*u*) |*i* Firing〉 of attractor patterns *s*(*u*) occurring whenever the landmark cell *i* fires can be written as

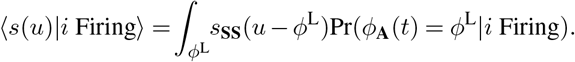

Thus all that matters for determining synaptic strength is the distribution of attractor phases that occur when the landmark cell fires.

Now, because the learning rule is linear and the landmark cell synapses only observe attractor steady states, the Hebbian weights W_*i*_ (*u*) can be written as a weighted superposition of the attractor bump patterns with weighting coefficients 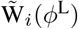:

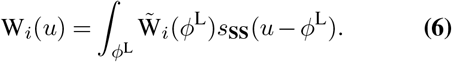

Furthermore, inserting Eq. (6) into Eq. (5) yields the learning dynamics of the synaptic weighting coefficients (see App. A for a proof):

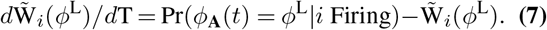

### Combined neural and synaptic dynamics during exploration

By combining the effect of path integration on the attractor phase *ϕ*_**A**_ described in Eq. (3) with the effect of multiple landmark cells with arbitrary learned weights 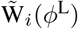, each acting on the phase through Eq. (4), we obtain the full dynamics of attractor phase driven by both animal velocity and landmark encounters:

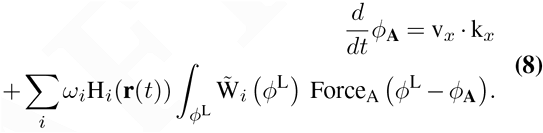

Together, Eq. 7 and Eq. 8 reflect a complex coupled dynamics between neurons and synapses. In Eq. 7 the distribution of attractor network activity patterns, or phases, drives plasticity in synapses from landmark cells to the attractor network. In turn, these synaptic weights modify the evolution of the attractor network phase via Eq. 8.

### Coupled landmark and attractor phase dynamics in the linearized model

The learned weights of a landmark cell are composed of a distribution of attractor network states. Linearizing Force_A_(*ϕ*^L^ − *ϕ*_**A**_) ≈ (*ϕ*^L^ − *ϕ*_**A**_), we can simplify this representation to a single variable: the weighted average of the distribution 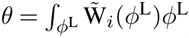, yielding a simplified equation for describing the neuronal and synaptic outcome of navigation and learning (see App. B):

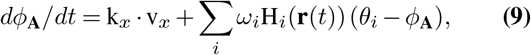

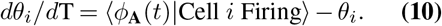

In essence Eq. 9 and Eq. 10 constitute a significant model reduction of Eq. 1 and Eq. 5. In this reduction, the entire pattern of neural activity of the attractor network is summarized by a single number *ϕ*_**A**_, denoting a point, or phase, on the ring manifold of stable attractor states. Similarly, the entire pattern of synaptic weights W_*i*_(*u*) from landmark cell *i* into the attractor network is summarized by a single number *θ_i_*, denoting the learned attractor network phase associated with the landmark cell’s synapses. Intuitively, the reduced Eq. 9 describes both path integration and a dynamics whereby each landmark cell *i* attempts to *pin* the attractor phase *ϕ*_**A**_ to the landmark cell’s learned phase *θ*_*i*_, each time the physical position **r**(*t*) of the animal is within the landmark’s firing field **H**_*i*_. In turn, synaptic plasticity described in Eq. 10 aligns the learned pinning phase *θ*_*i*_ of each landmark cell *i* to the average of the ensemble of attractor phases *ϕ*_**A**_ that occur when the animal is in the firing field of the landmark.

As we will see below, as an animal explores its environment, this coupled dynamics between attractor phase *ϕ*_**A**_ and landmark pinning phases *θ*_*i*_ settle into a self-consistent steady state such that the attractor phase yields an internal estimate of the animal’s current position that is, to first order, largely independent of the history of the animal’s previous trajectory. Moreover, each landmark cell learns a pinning phase *θ*_*i*_, consistent with the location of its firing field in physical space.

## Learning a simple environmental geometry

We now examine solutions to these equations to understand how neuronal dynamics and synaptic plasticity conspire to yield a consistent map of the environment. To build intuition, first consider the linearized dynamics of Eqs. 9, 10 for the simple case of an animal moving back and forth between the walls of a 1D box of length L, at a constant speed v = L/*τ*, yielding a total time of 2τ to complete a full cycle (Fig. 3A). In this environment we assume two landmark cells corresponding to the east (west) walls, with firing fields extending a distance **L_Wall_** into the environment leaving an empty space **L_Int_** = **L** − **2L**_**Wall**_ between. Their pinning phases *θ*_East_ (*θ*_West_) encode the peak position of their outgoing synaptic weights. How does circuit plasticity yield a consistent environmental representation through exploration?

**Fig. 3.**
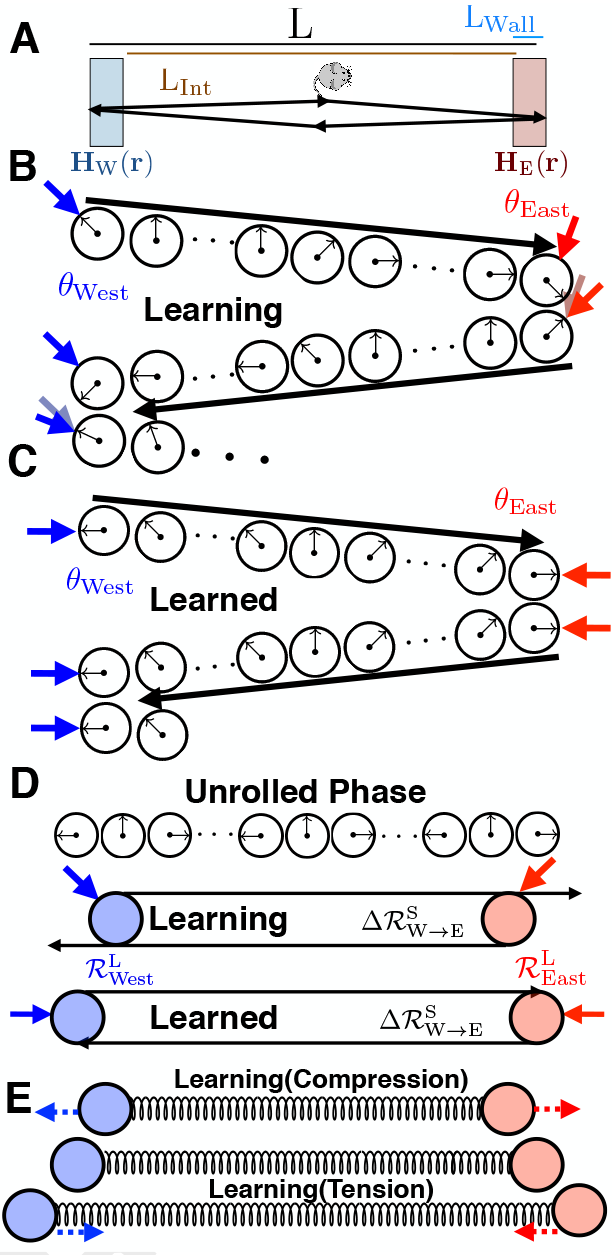
**A)** An animal moving between two landmarks at the edges of a 1D track. **B)** A single cycle of exploration as the animal moves from the west to east wall and back. When the animal encounters the west (east) wall, the attractor phase (black arrow) is pinned to the associated landmark pinning phase (blue/red arrow for west/east wall). As the animal moves from one wall to the other, the attractor phase advances from this pinned phase due to path integration. During learning, the pinning phase from any one wall, plus the phase advance due to path integration, will not equal the pinning phase of the other wall. However, plasticity will adjust the pinning phase of each wall to reduce this discrepancy (motion of red and blue arrows). During this inconsistent pre-learned state, the attractor phase at any interior position will depend on path history. **C)** After learning, the pinning phase from any one wall, plus the phase advance due to path integration, *equals* the pinning phase of the other wall, yielding a consistent internal representation of space in which the attractor phase assigned to any interior point becomes independent of path history. **D)** We can “unroll” the attractor and landmark phases into linear position variables. Thus landmark cell synapses can be thought of as points in physical space (blue and red circles). If the phase advance due to path integration *exceeds* phase difference between the pinning phases of the landmarks, then the distance between the landmark cells in unrolled phase is *closer* than the physical distance between the firing fields of the landmarks (top). Plasticity then exerts an outward force pushing the two landmark cells further apart until their separation in unrolled phase equals the physical distance between between their firing fields (bottom). **E)** In general, the changing positions in unrolled phase associated with landmark cell synapses due to synaptic plasticity can be described by a damped springlike interaction as in Eq. 11 and Eq. 12. If the separation between the two landmark cell synapses in unrolled phase is smaller (larger) than the physical separation between their firing fields, then the spring will be compressed (extended), yielding an outward (inward) force. This force will move the positions associated with landmark cell synapses until their separation in unrolled phase equals the rest length of the spring, which in turn equals the physical separation between landmark firing fields.

We will build intuition in the limit where **L**_**Wall**_ → 0, *ω* → ∞; in this regime, landmark cells only act at the very edge, yet *fully* anchor the attractor state when the animal touches the edge. At *t* = 0, the animal starts at the west wall at physical position **r**(0) = −L/2. Through Eq. 9, the west border cell pins the initial attractor phase so that *ϕ*_**A**_ (0) = *θ*_West_. At *t* = *τ*, the animal travels to the east wall at physical position **r**(*τ*) = +L/2, and the attractor phase advances due to path integration to become to become *ϕ*_**A**_(*τ*^−^) = *θ*_West_ + k_*x*_L. However, upon encountering the east wall, the east border cell pins the attractor phase to *θ*_East_.

Before any learning, there is no guarantee that the east border cell pinning phase *θ*_East_ equals the attractor phase *θ*_West_ + k_*x*_L, obtained by starting at the west wall and moving to the east wall; sensation and path integration might disagree (Fig. 3B). However, plasticity described in Eq. 10 will act so as to move *θ*_East_ closer to *θ*_West_ + k_*x*_L. Then as the animal returns to the left wall at time *t* = *2τ*, path integration will retard the attractor phase *ϕ*_**A**_(2*τ*) = *θ*_East_ − k_*x*_L, and an encounter with the west wall leads the west border cell to pin the attractor phase to *θ*_West_. Again, there is no guarantee that the west border cell pinning phase *θ*_West_ agrees with the attractor phase *θ*_East_ − k_*x*_L obtained by starting at the east wall and traveling to the west wall, but circuit plasticity will change *θ*_West_ to reduce this discrepancy. Overall, plasticity over multiple cycles of exploration yields the iterative dynamics

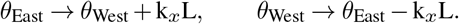

Thus the phase difference *θ*_East_ − *θ*_West_ between the pinning phases of the two landmark cells will approach the phase advance k_*x*_L incurred by path integration between the two landmarks. Thus learning can precisely co-register sensation and path integration so that these two information sources yield a consistent map of space (Fig. 3C). In particular, the attractor phase assigned by the composite circuit to any point in the interior of the environment now becomes independent of which direction the animal is traveling, in contrast to the case before learning (compare the assigned interior phases in Fig. 3B versus C).

### Learning as an elastic relaxation between landmarks

To gain further insight into the learning dynamics, it is useful to interpret the periodic attractor phase *ϕ*_**A**_(*t*) as an internal estimate of position through the “unrolled” coordinate variable 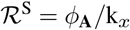. Likewise, we can replace the landmark phase *θ*_*i*_ with another linear variable RL = 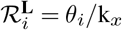, denoting the internal representation of the position of landmark *i* (Fig. 3D). This enables us to associate physical positions to landmark cells, or more precisely their pinning phases, although these assigned positions are defined only up to shifts of the grid period. Plasticity over the long timescale T of exploration then yields the following learning dynamics for the physical positions in unrolled phase for the landmark cells:

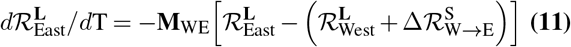

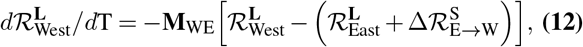

where 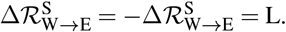

This dynamics for the two landmark cell synapses in unrolled phase is equivalent to that of two particles at physical positions 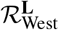 and 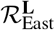, connected by an overdamped spring with rest length L, and spring constant **M**_WE_ which sets the learning rate (Fig. 3E). If the separation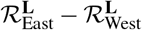 between the particles is less (greater) than L, then the spring is compressed (extended) yielding a repulsive (attractive) force between the two particles. Learning stabilizes the two particle positions when their separation equals the spring rest length, so that 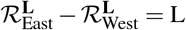. This condition in unrolled phase is equivalent to the fundamental consistency condition for a well defined spatial map, namely that the phase advance due to path integration equals the phase difference between the pinning phases of landmark cells (Fig. 3C). However the utility of the unrolled phase representation lies in revealing a compelling picture for how a spatially consistent map arises from the combined neuronal and synaptic dynamics, through a simple, emergent first order relaxational dynamics of landmark particles connected by damped springs. As we see below, this simple effective particle-spring description of synaptic plasticity in response to spatial exploration generalizes to arbitrary landmarks in arbitrary two dimensional environments.

We note that if the environment has not been fully learned or has been recently deformed, the internal representation of landmarks in unrolled phase will lag behind the true geometry for a time, leading to “boundary-tethered” firing fields seen in (22, 30). Additionally, we have solved the dynamics when the firing fields of the border cells have a finite extent L_**Wall**_ and the landmark cells have a finite strength *ω*, and we find the dynamics obeys that of Eq. 11 and Eq. 12, with a different rest length 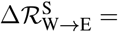 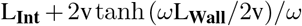 (See App. D).

## Generalization to 2D Grid Cells

In order to make contact with experiments, we generalize all of the above to two dimensional space. Now grid cells live on a periodic *two-dimensional* neural sheet, where each neuron has position **u** = (*u*_1_,*u*_2_). The dynamics, analogous to Eq. 1 are:

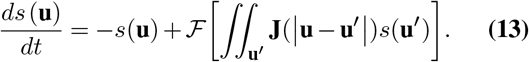

Where **J** includes short-range excitation and long-range inhibition(Fig. 4A). Attractor dynamics on a *two-dimensional* neural sheet can now yield a *two-dimensional* family of stable, or steady state, localized bump activity patterns 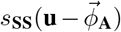 with *hexagonal* symmetry (App. E). The attractor state is now a 2D phase 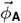 on the periodic rhombus (Fig. 4B).

**Fig. 4.**
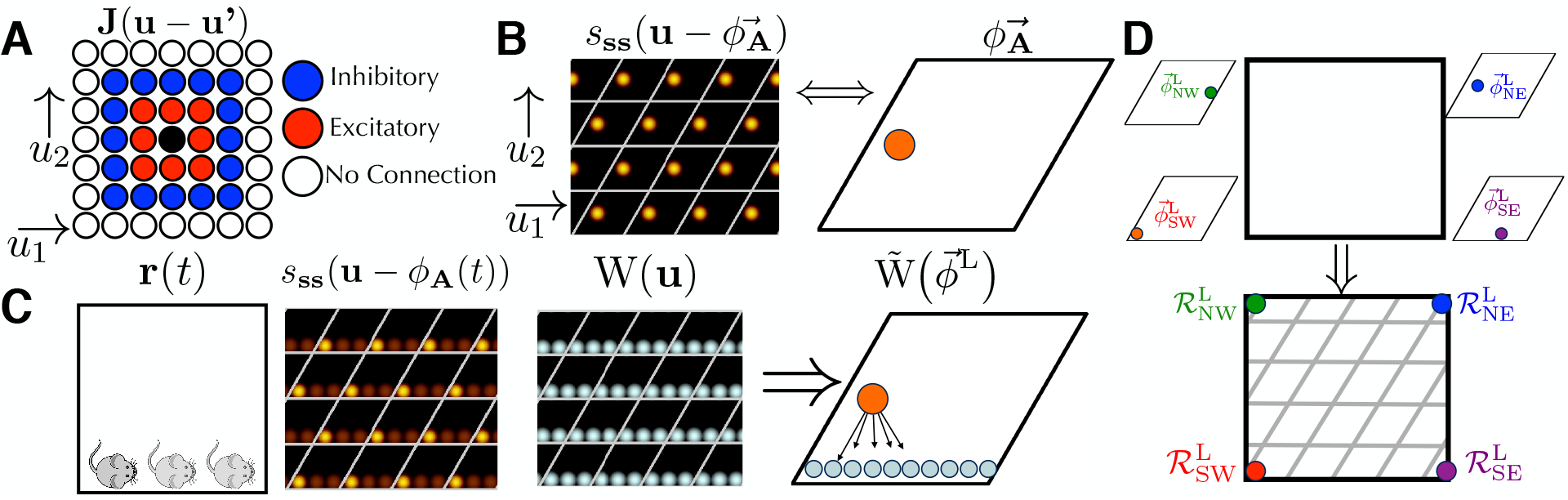
**A)** A 2D neural sheet with short-range excitation and long-range inhibition, analogous to Fig. 1. Each neuron on the continuous sheet now has coordinates **u** = (*u*_1_,*u*_2_). **B)** A 2D analogue of a single attractor pattern on the neural sheet, with high firing rates in red (compare to Fig. 1). The set all unique stable attractor patterns is now indexed not by a single phase variable as in 1D, but a 2D phase variable 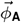 A ranging over a rhombus, or unit cell. Copies of the unit cell are shown via white lines. **C)** The landmark cell hebbian weights will be a combination of 2D attractor states (Eq. 15). As the animal travels along the south wall, the average firing rates will form a “streak” across the neural sheet. This leads the hebbian weights on the neural sheet to form the same streak; this learned state can be represented as a distribution over the periodic rhombus. Analogously, there is a force law, where the state of an attractor network 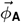 will be pulled towards this distribution 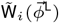 (Eq. 14). D) Similarly to Fig. 3D, we can unroll the two-dimensional attractor phase into a two-dimensional position variable, thereby associating landmark pinning phases to points in physical space. Given landmarks in all four corners, the landmark pinning phases correspond to different points on the phase rhombus, but through unrolling this rhombus, each can be associated to a physical corner of the environment.

Applying the same techniques used to derive Eq. 8, we obtain a 2D analogue to dynamics of the attractor state:

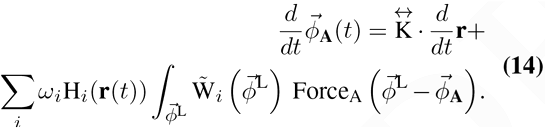

Here we replaced k_*x*_ in 1D with 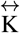, a matrix that translates 2D animal velocity into phase advance in the 2D attractor network; 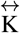 determines *both* grid spacing *and* orientation. The analog of learning dynamics in (Eq. 7) is:

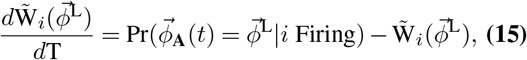

where 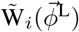 is now a distribution over the periodic rhombus (Fig. 4C).

In an analogous manner, we may make a small-angle approximation to replace the attractor phase *ϕ*_**A**_ (*t*) with a *two-dimensional* attractor coordinate variable 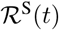, reflecting an internal estimate of instantaneous position in physical space, and we replace the landmark phase 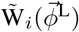 with another 2D attractor coordinate variable 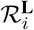 reflecting the internal estimate of landmark position(Fig. 4D). This yields two-dimensional dynamics for position self-estimates and landmark position, given in analogy to Eqs. 9, 10 by:

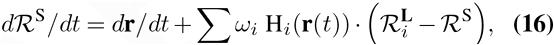

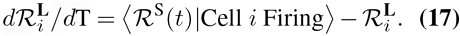

## Spatial consistency through emergent elasticity

We showed in Eq. 11 and Eq. 12, and in Fig. 3DE that the emergence of spatial consistency between path integration and landmarks through Hebbian learning dynamics, during exploration of a simple 1D environment, could be understood as the outcome of an elastic relaxation process between landmark cell synapses, viewed as particles in physical space connected by damped springs. Remarkably, this result generalizes far beyond this simple environment. As long as the exploration dynamics are time-reversible^1^, the learning dynamics of *any* set of landmark cells in *any* geometry yields this particle-spring interpretation:

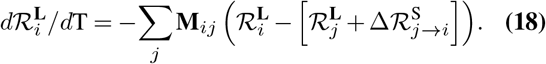

The spring constant **M**_*ij*_ is related to the frequency with which the animal moves between each pair of landmark firing fields *i*, *j*, while the rest displacement 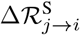 is the average change in unrolled attractor phase as the animal moves from firing field *j* to field *i*, roughly related to the distance between the landmark firing fields. Precise expressions for the spring constants and rest lengths are derived from the statistics of exploration in (App. F).

Overall, this elastic relaxation process converges towards an internal map where all pairs of landmark cell synapses, viewed as particles in unrolled phase, or physical space, become separated by the physical distance between their firing fields. This convergence ensures a consistent internal environmental map of external space in which velocity based path integration of attractor phase starting at the pinning phase of landmark *i* and ending at landmark *j* will yield an integrated phase consistent with the pinning phase of landmark *j* itself. This relaxation dynamics explains path-dependent shifts in firing patterns observed in recently deformed environments (22). Also, the experimental observation that in multi-compartment environments, consistent maps *within* compartments form before consistent maps *between* compartments are also explained (10) by this relaxation dynamics. In essence, the longest-lived learning mode of the relaxation dynamics corresponds to differences in maps *between* compartments.

Furthermore, as we explain in the next three sections, these relaxation dynamics yields several novel experimental predictions: (1) systematic path-dependent shifts in *fully learned* 2D environments, (2) mechanical deformations in complex environments, and (3) the novel prediction of creation of topological defects in grid cell firing patterns through specific environmental manipulations.

## Path-dependence in 2D environments

We saw above that exploration in a simple 1D geometry lead to a consistent internal map in which the attractor network phase was mapped onto the current physical position alone, independent of path history (Fig. 3C). This consistency arises through the elastic relaxation process in Eq. 11 and Eq. 12, which makes the distance between the landmark cells in unrolled phase 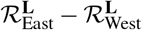 equal to the physical distance between their firing fields L, just like two particles connected by a spring with rest length L (Fig. 3E). This situation will generalize to two dimensions if there are only two landmarks, namely a west and east border cell (Fig. 5A1). However, it becomes more complex with the addition of a third landmark cell, for example a south border cell (Fig. 5A2).

**Fig. 5.**
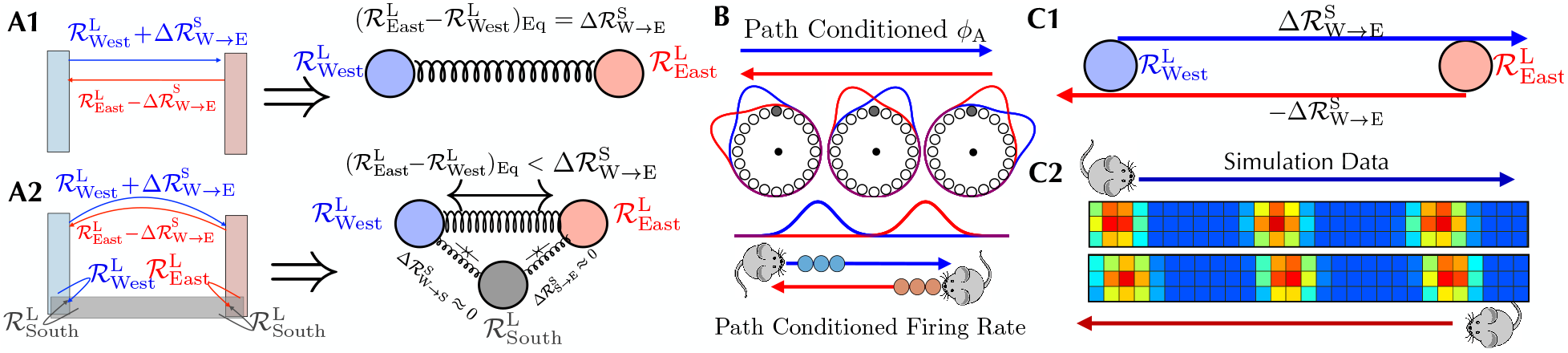
**A1)** For two landmark cells, the rest length 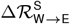 of the spring connecting them equals the physical width L of the environment, and so the two landmark particles learn unrolled pinning phases 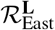 and 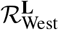 obeying the spatial consistency condition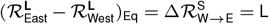 as in Fig. 3C. **A2)** The addition of a southern landmark cell will cause a pinning effect which pulls 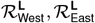 closer together. The animal can travel from the east and west landmark field to the southern landmark field with little path integration at all, yielding 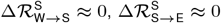. **B)** If the attractor phase is advanced on a west to east trajectory (blue) relative to an east to west trajectory (red), then any particular grid cell (in this case the shaded grey cell) will fire earlier (later) on west-to-east (east-to-west) trajectory. Thus grid fields computed from trajectories leaving the west (east) border will shifted west (east). **C1)** When landmark pinning phases are pulled together closer than the path integration distance between them, then the attractor phase will shift *away* from whichever wall the animal last encountered. Therefore it will phase advance on west-to-east trajectories relative to east-to-west trajectories, as in Fig. 3B and Fig. 5B. **C2)** Thus simulations of Eq. 14 and Eq. 15 lead to grid cell firing patterns shifted *towards* whichever wall the animal last encountered.

In this case, east and west landmark particles will be connected by a spring of rest length 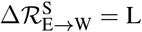, as before, but they will each also be connected to the south landmark particle with springs. Intuitively, as the mouse travels from the east or west walls to the south walls, the landmark pinning phases of each of these three border cells will be attracted towards each other^2^. The combined three particle elastic system will settle into an equilibrium configuration in which the difference in unrolled phase between east and west landmarks will be *less* than the physical separation L, or equivalently the rest length 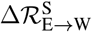 of the spring connecting them. This in turn implies that the attractor phase assigned to any physical position in the interior will be relatively phase advanced (retarded) if the mouse is on a trajectory leaving the west (east) wall. This path dependence in the attractor phase is entirely analogous to that seen in Fig. 3B. However, the reason is completely different. In Fig. 3B, the landmark particles are not separated by the rest length of the spring connecting them because the environment is not fully learned and so the particles are out of equilibrium, whereas in Fig. 5A2, the particles are not separated by the rest length, even in a *fully* learned environment, because additional springs from the south landmark create excess compression.

This theory makes a striking experimentally testable prediction, namely that even in a *fully learned* 2D environment, grid cell firing fields, when computed on subsets of mouse trajectories conditioned on leaving a particular border, will be shifted *towards* that border (Fig. 5B). This shift occurs because at any given position, the attractor phase depends on the most recently encountered landmark. In particular, on a west to east (east to west) trajectory, the attractor phase will be advanced (retarded) relative to a east to west (west to east) trajectory. Thus on a west to east trajectory, the advanced phase will cause grid cells to fire earlier, yielding west shifted grid cell firing fields as a function of position. Similarly on an east to west trajectory, grid fields will be east shifted. In summary, the theory predicts grid cell firing patterns conditioned on trajectories leaving the west (east) border will be shifted west (east). While we have derived this prediction qualitatively using the conceptual mass-spring picture in Fig. 5A2, we confirm this intuition through direct numerical simulations of the full circuit dynamics in Eq. 14 and Eq. 15 (Fig. 5C2). Under reasonable parameters, our simulations can yield path-dependent shifts of up to ~2 cm towards whichever wall the animal last touched (App. G).

We searched for such subtle shifts in a population of 143 grid cells from 14 different mice that had been exploring a familiar, well-learned, 1-meter open field (App. H), using two separate analyses, based on cross-correlations and spike shifts with respect to field centers.

### Cross-Correlations

One method for detecting a systematic firing field shift across many grid fields is to cross-correlate firing rate maps conditioned on trajectories leaving two different borders (App. A.1). For example, fer euch cell, we can ask how much and in what direction we must shift itn west border conditioned firing field to match, or correlate as much as possible with, the same cell’s east conditioned firing field. In particular, for each cell, we can compute the correlation coefficient between a spatially shifted west conditioned field and an unshifted east conditioned field, and plot the average correlation coefficient as a function of this spatial shift. The theory predicts that we will have to shift the west conditioned firing field eastwards to match the east conditioned firing field (Fig. 6A). This prediction is confirmed by a peak in the cross-correlation as a function of spatial shift when the spatial shift is positive, or directed east (Fig. 6B).

**Fig. 6.**
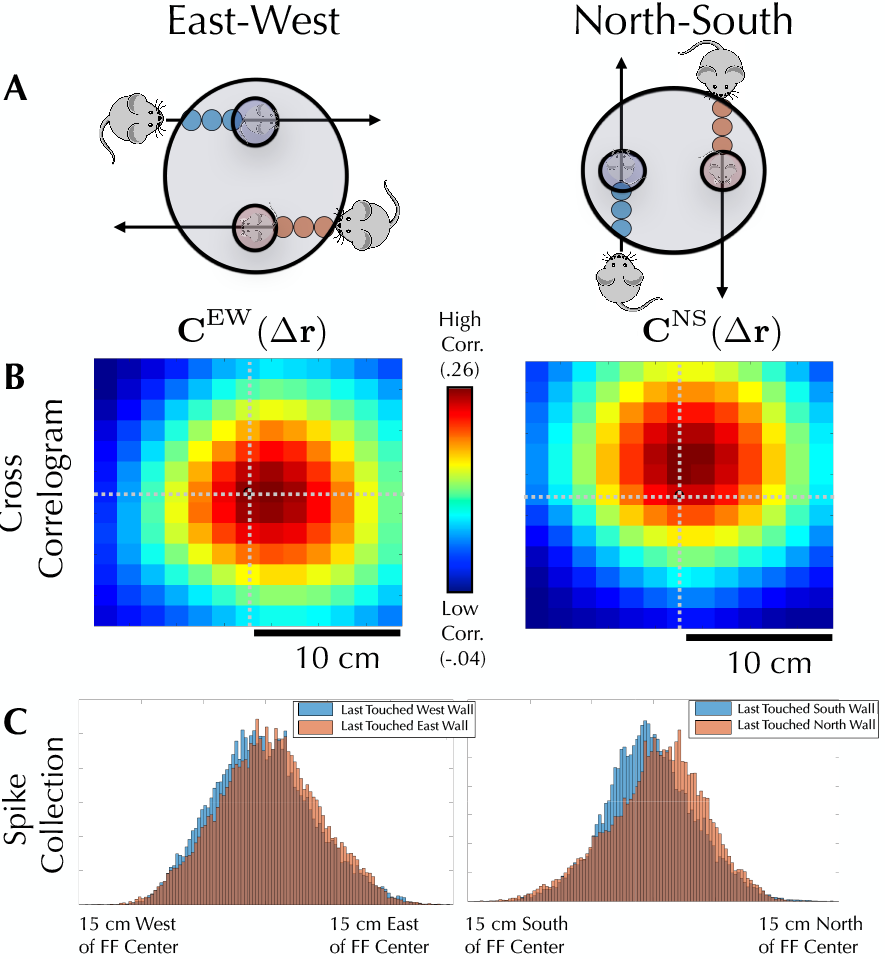
**A)** Our theory predicts that grid cell firing patterns will be shifted towards whichever wall the animal last encountered, even in a fully learned environment. **B)** On the left (right) this shift is detected by computing the cross correlation between west (south) conditioned firing fields, shifted by a spatial offset Δr, and the unshifted east (north) conditioned firing field. The cross correlation peaks when the spatial shift is positive in the x-direction (positive in the y-direction), as predicted by theory. **C)** This effect can also be seen by comparing histograms of spike positions around firing field centers for different path conditions.

A similar logic holds for north and south; in order to maximally correlate the south conditioned field to the north conditioned field, the theory predicts we will need to shift the south conditioned map north. This requisite shift is seen in the data in Fig. 6B, which reflects the cross correlation between a shifted south field and an unshifted north field, averaged across all cells. The maximal correlation is achieved when the south fields are shifted north. Overall, this analysis shows that grid patterns are shifted towards the most recently encountered wall, both for the NS walls (3 cm, *P* = 1.5 · 10^−5^, Binomial Test, *P* = 1.5 · 10^−5^, Sign-Flip Test) and the EW Walls (1.5 cm, *P* = 10^−7^, Binomial Test, *P* = 10^−7^, Sign-Flip Test), matching the sign and magnitude seen in simulations.

### Firing Field Centers

These results can be corroborated by computing shifts in spikes relative to firing field centers, when conditioning spikes on the path history (App. A.2). For each firing field center, we calculate the average spike position within that firing field conditioned on the animal having last touched a particular wall. For each cell, we calculate the average shift across all firing fields, and examine how the shifts depend on which wall the animal last touched. Again, the patterns are shifted towards whichever wall the animal last touched(Fig. 6C) for both the NS walls (.5 cm, *P* = 10^−5^ Binomial Test, *P* = 10^−5^ Sign-Flip Test) and the EW Walls (.5 cm, *P* = 3 · 10^−4^ Binomial Test, *P* = 2 · 10^−2^ Sign-Flip Test). The discrepancy in the estimated magnitude of the shift between the methods of analysis is likely due to poorly defined firing fields; a method based on firing field centers will give a lower signal-to-noise ratio, and thus a lower shift magnitude, than the cross-correlogram method.

## Mechanical deformations in complex environments

Another experimental observation that can be reproduced by our theory is the distortion (11) of grid cell patterns seen in an irregular environment(Fig. 7A). Landmark cells with firing fields distributed across an entire wall will pull the attractor phase to its associated landmark pinning phase *regardless* of where along the wall the animal is. The presence of a diagonal wall then causes the average attractor phase as a function of position to curve towards the wall, yielding spatial grid cell patterns that curve *away* from the wall (Fig. 7B, C). Previous theoretical accounts of this grid cell deformation have relied on purely phenomenological models that treated individual grid cell firing fields as particles with mostly repulsive interactions (15), without a clear mechanistic basis underlying this interaction. Here we provide, to our knowledge for the first time, a model with a clear mechanistic basis for such deformations, grounded in the interaction between attractor based path integration and landmark cells with plastic synapses. Such dynamics yields an emergent elasticity where the particles are landmark cell synapses rather than individual firing field centers.

**Fig. 7.**
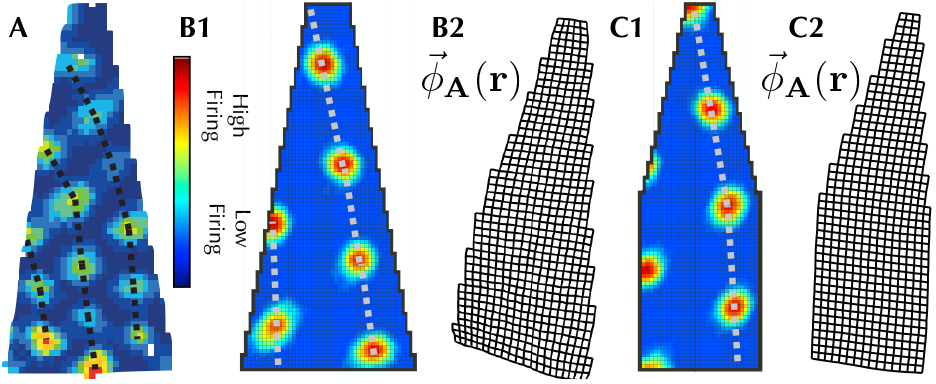
**A)** Experimental data of grid cell firing patterns deformed, curving *away* from a wall in an irregular geometry. **B1)** A full simulation of Eq. **14**, Eq. **15** also yields grid firing patterns bent away from the wall. **B2)** Visualization of the average attractor state as a function of position 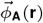(periodicity removed for visualization purposes). The reversal between the bending of the internal attractor phase and the bending of firing rate maps is similar to the reversal seen in Fig. 5B. **C1)**, **C2)** Same as B1), B2), but for a slightly different geometry.

## Topological defects in grid cells: a prediction

While the dynamics of the linearized Eq. 17 will always flow to the same relative landmark representations 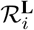, this is not the case for the full dynamics of Eq. 14, Eq. 15, which can learn multiple different stable landmark cell synaptic configurations. One striking example of this is the ability of the learning dynamics to generate “topological defects”, where the number of firing fields traversed is not the same for two different paths (Fig. 8A, B and App. G). An environmental geometry capable of supporting these defects will yield a set of firing patterns that depends not only on the final geometry, but also on the *history* of how this geometry was created (Fig. 8C).

**Fig. 8.**
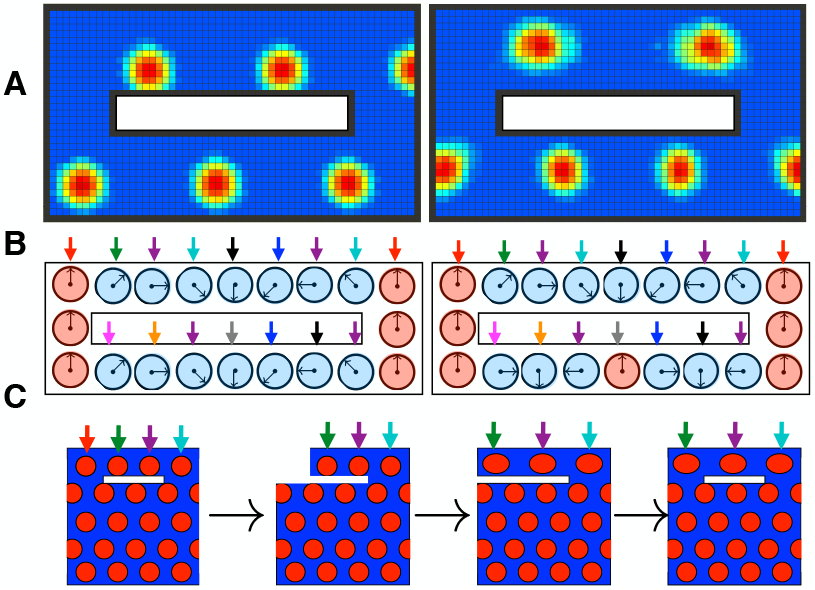
**A)** Two steady state grid cell patterns emerging from the same cue-rich environment. In the first firing pattern, the combination of landmark pinning and path integration yields a phase advance of four firing fields in traveling from west to east along either corridor. The second pattern has a topological defect; traveling from the west to east through the *north* corridor yields a phase increase of ~ 1.5 firing fields; traveling east to west through the *south* corridor yields a phase decrease of ~ 2.5 firing fields. This second pattern is stable nonetheless. **B)** Schematic of 1D underlying attractor state as a function of space. The two patterns in (A) correspond to two different landmark pinning phase patterns learned by the many landmarks. Both landmark pinning patterns are stable under Eq. 14, Eq. 15. In the first pattern, the combination of landmark pinning and path integration yields the same phase advance in both the north and south corridors. The second pattern has a topological defect; the phase advance in the north corridor *one full rotation less* than the phase advance through the south corridor. This is possible because many landmark cues (colored arrows) can yield many landmark cells with multiple stable synaptic configurations, or pinning phases under Eq. 14, Eq. 15. **C)** Schematic of proposed “deformation schedule” that could yield a topological defect in grid cell firing patterns. By separating/truncating the northern corridor, stretching it (along with spatial cues, denoted by colored arrows), than reconnecting it, it may be possible introduce one of these defects. Even though the initial geometry is identical to the final geometry, the deformation schedule has lead to a firing pattern which is three fields wide in the north and four fields wide in the south.

## Discussion

Overall, we have provided a theoretical framework for exploring how sensory cues and path integration may work together to create a consistent internal representation of space. Our framework is grounded in biologically plausible mechanisms involving attractor based path integration of velocity and Hebbian plasticity of landmark cells. Moreover, systematic model reduction of this combined neural and synaptic dynamics yields a simple and intuitive emergent elasticity model in which landmark cell synapses act like particles sitting in physical space connected by damped springs whose rest length is equal to the physical distance between landmark firing fields. This simple emergent elasticity model not only provides a conceptual explanation of how neuronal dynamics and synaptic plasticity can conspire to self-organize a consistent map of space in which sensory cues and path-integration are in register, but also provides novel predictions involving small shifts in firing fields even in fully learned environments, the possibility of topological defects in grid cells, and the mechanical deformation of grid cells in response to irregular borders.

This work opens up many interesting avenues for future research. For example, further explorations of the nonlinear regime of our combined circuit dynamics may yield interesting experimental signatures that distinguish different modes of interactions between attractor networks, path integrators and landmark cells. Incorporating heterogeneity of neural representations observed in MEC (31) into our framework is another intriguing avenue. Also, as the reliability of sensory and velocity cues change, it is interesting to ask what higher order mechanisms may exist to differentially regulate the effect of landmarks and velocity on the internal representation of space. More generally, our theory provides a unified framework for understanding how systematic variations in environmental geometry and the statistics of environmental exploration interact to precisely sculpt neural representations of space.

## Acknowledgements

LMG is a New York Stem Cell Foundation - Robertson Investigator. This work was supported by funding from The New York Stem Cell Foundation, James S McDonnell Foundation, Whitehall Foundation, NIMH MH106475 and a Klingenstein-Simons Fellowship awarded to LMG, funding from the Simons Foundation awarded to LMG and SG. SO was supported by the Karel Urbanek Postdoctoral Fellowship in Applied Physics. KH is supported by a Stanford SIGF. We thank Daniel Fisher for helpful discussions.

## Supplementary Note A: Reducing perturbation to an effective force law in the one-variable “ring” representation

Consider a steady-state firing rate pattern *s*_**SS**_(*u* − *ϕ*_**A**_) under some translation-invariant neural dynamics function Dyn that is given some perturbation centered at *ϕ*^**p**^ having the same periodicity as *s*_**SS**_:

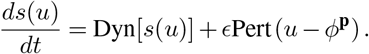

In order to understand the effect of this perturbation, we need to understand the Jacobian matrix around the point *s*_SS_ (*u* − *ϕ*_**A**_):

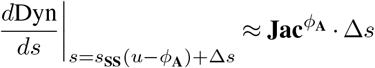

### A. Modes of the Jacobian

Because *s*_**SS**_ (*u* − *ϕ*_**A**_) is a stable one-dimensional family of solutions of Dyn,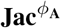 must be a negative semidefinite matrix. Because

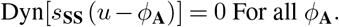

there is a single-zero eigenvector^3^, the sliding mode 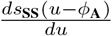:

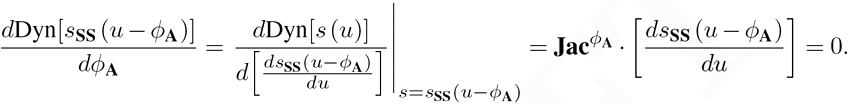

### B. Effect of small perturbations

When an external perturbation is small and 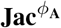 is symmetric, e.g. 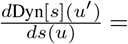 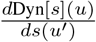, the effective perturbation will the the projection of the actual perturbation onto the sliding mode.

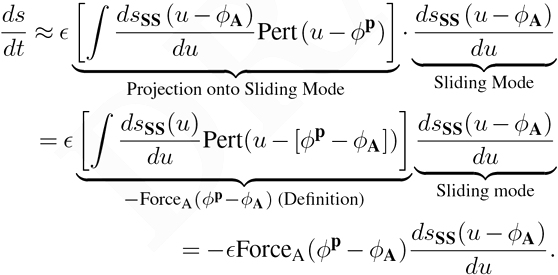

We can translate these dynamics into the reduced *ϕ* representation:

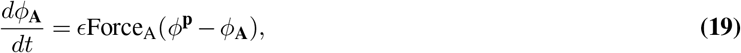

Eq. 19 can be verified:

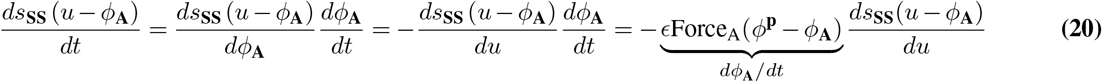

#### B.1. Form of the landmark cell force function

When the perturbation takes the form of input from Hebbian landmark cells, the perturbation function is simply the attractor bump pattern *s*_**SS**_(*u* − *ϕ*^L^). Therefore, defining Δ*ϕ* = *ϕ*^L^ − *ϕ*_**A**_,

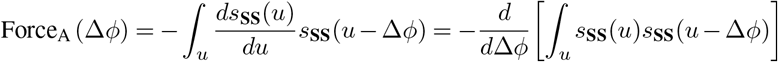

Therefore, the force function is simply the negative derivative of the spatial autocorrelation function of the bump pattern. Because the spatial autocorrelation is even and maximized at Δ*ϕ* = 0, minimized at Δ*ϕ* = π, the force function will be odd, with positive(negative) values for positive(negative) Δ*ϕ*. As long as the bump size is not much smaller than the bump spacing, the autocorrelation will decrease gradually between Δ*ϕ* = 0, Δ*ϕ* = π, leading to a long range force function which only approaches zero at, and far from, the origin. This behavior is qualitatively matched by Force_A_ (Δ*ϕ*) = sin(Δ*ϕ*). For simplicity, we define the magnitude of Force_A_(Δ*ϕ*) to give it a slope of 1 at Δ*ϕ* = 0; all strength information can be contained in *ω*.

#### B.2. Dynamics with non-symmetric Jacobians

When 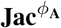 is non-symmetric, we may use the same techniques as before, except now we must use a non-orthogonal projection onto the sliding mode:
Non-orthogonal projection onto sliding mode:

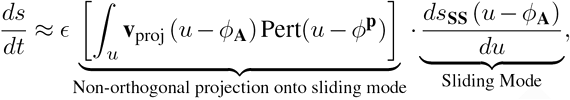

where **v**_proj_(*u* − *ϕ*_**A**_) can, in principle, be solved through diagonalization of the Jacobian 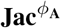.

## Supplementary Note B: Proof of recovery of exact path integration

By coupling the attractor network to conjunctive position and velocity-tuned cells that east (west) movement-selective cells form feedforward synapses into the attractor network that are shifted in the positive (negative) *u* (28)direction, we impose a velocity-dependent perturbation on the network. This yields,

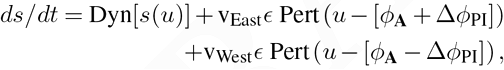

where v_East_, v_west_ are the east and west velocities of the animal. Then model reduction via Eq. 2 yields

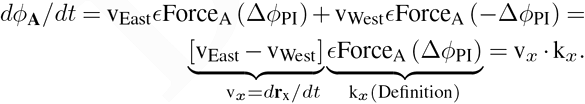

Here k_*x*_ is a constant of proportionality that relates animal velocity to the rate of phase advance in the attractor network.

## Supplementary Note C: Lemmas about Landmark Cells

### A. Verifying the Hebbian learning rule in the attractor basis

We can verify that in the attractor basis,

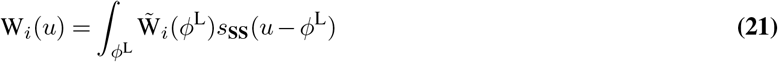

the learning rule:

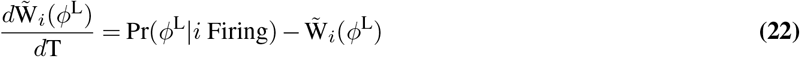

gives us the learning rule in the neural basis:

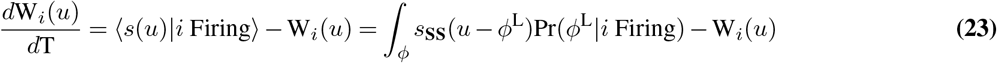

by inspection:

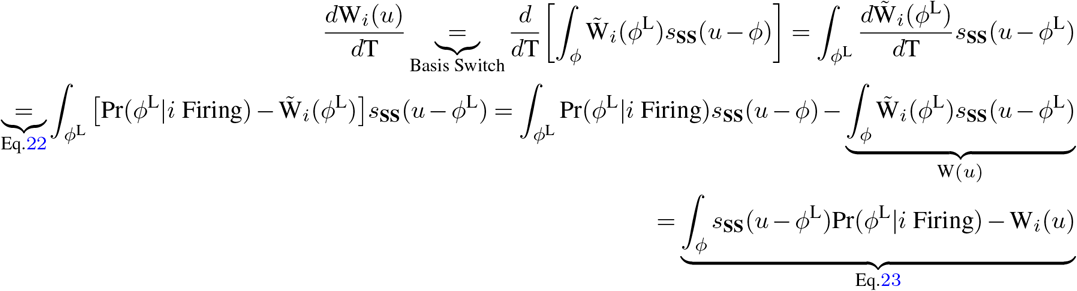

### B. Lemmas about linear approximations

#### B.1. Linear Approximation of Forcing Rule

We can show how linearizing the force law into a simple “spring constant” allows us to represent the landmark state with a single number 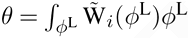:

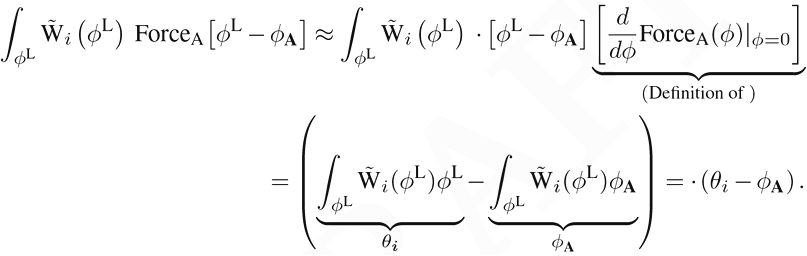

#### B.2. Proof of single-variable representation of landmarks

Combining the attractor-basis dynamics for hebbian learning and the linear approximation:

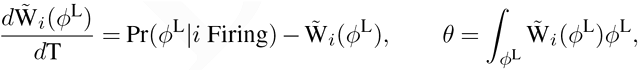

we get:

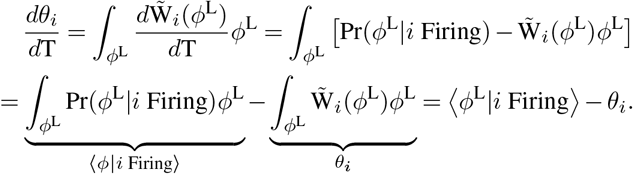

## Supplementary Note D: Proof of simplest case

The position self-estimate will reach a steady cycle, so we start with a animal at 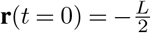, having position selfestimate 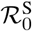. The position self-estimate will follow the linearized dynamics, which include terms for path integration aswell as the east and west landmarks.

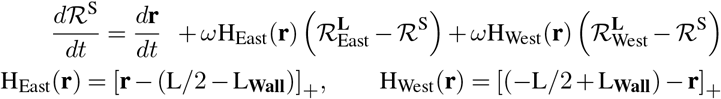

We can assume the position self-estimate will reach a steady cycle such that 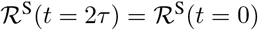. Defining 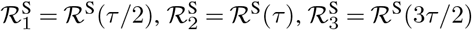, we can solve for the the position self-estimate as a piecewise function:

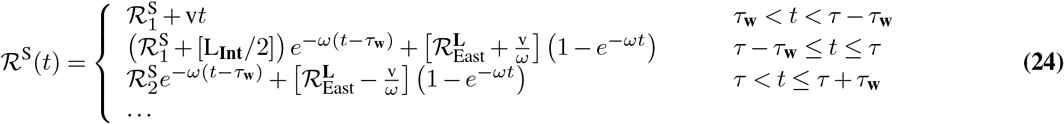

Where *τ*_**w**_ = **L_Wall_**/v. This yields a set of linear equations:

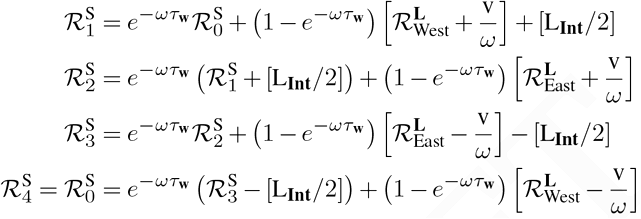

The average position self-estimate seen by the east landmark comes from two components of piecewise function 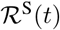. The first is *τ* − τ_w_ < *t* < *τ*, the second is *τ < t < τ* + τ_w_:

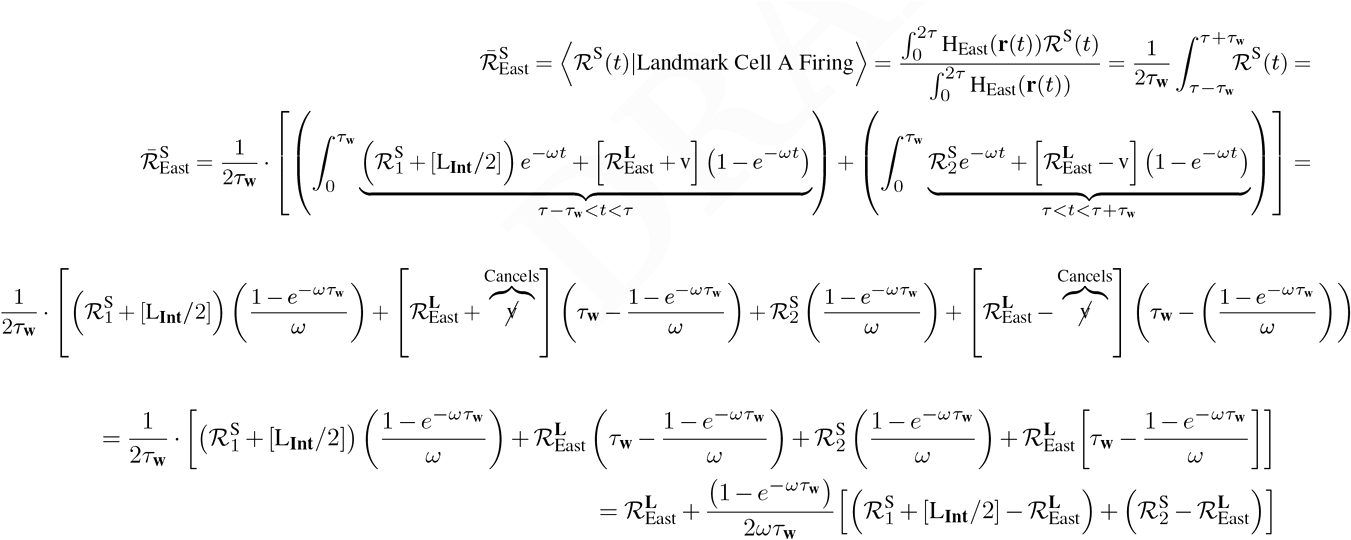

Therefore, at equilibrium:

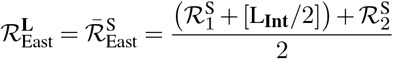

There is a translational symmetry to this problem, such that any shifted version of a solution is also a solution. We center around zero for simplicity, such that 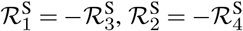 and 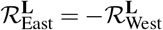. Combining the above equations and this symmetry gives the steady state solution:

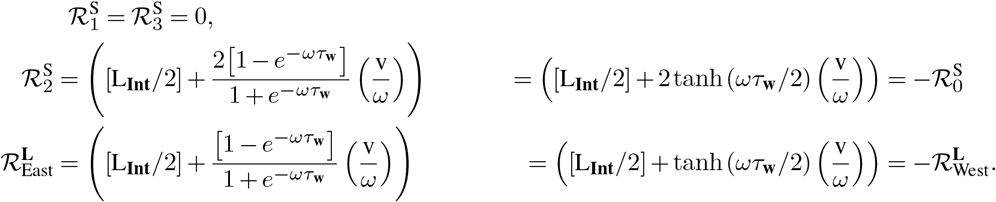

### A. Out of Equilibrium Path-Dependent Shifts and Learning Dynamics

When the system is out of equilibrium it is convenient to refer to the landmark representations in terms of their deviation from the equilibrium state.

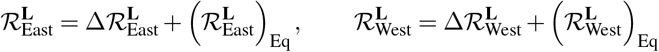

We have the set of linear equations for how much the position self-estimates vary with the landmark position estimates, where we use the shorthand 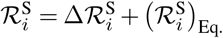:

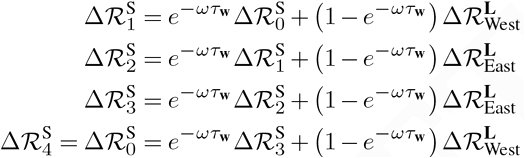

We can combine the equations for 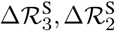 to get:

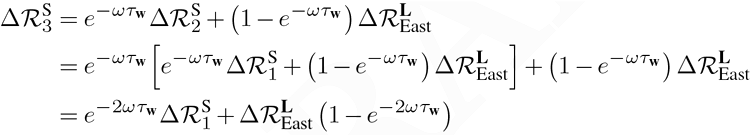

We can express through 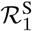 in terms of 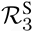 through symmetry:

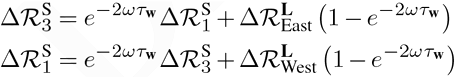

Plugging one into the other:

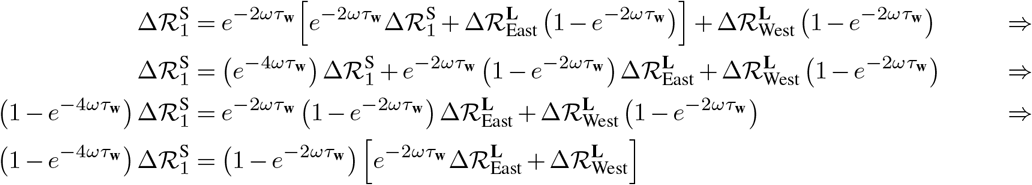

This yields the change in position self-estimate:

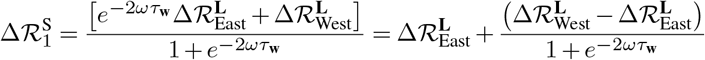

This allows us to recover the first coefficient related to path-dependent shift. When 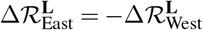

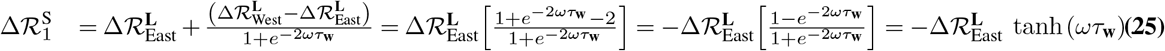

We note that when *ωτ*_**w**_ → ∞, the path-dependent shift is *exactly* the shift in the estimated position of the landmark last touched 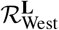. When *ωτ*_**w**_ → 0, the shift goes to 0, as the memory of 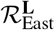 is nearly the same as that of 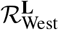.

#### A.1. Learning Timescale Coeffient

In order to understand the learning dynamics, we must calculate the effect of the *estimated* landmark position on the *estimated* position self-estimate that becomes associated with each landmark:

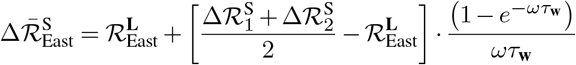

Plugging in:

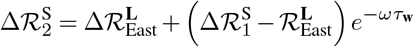

Gives things in terms of 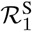:

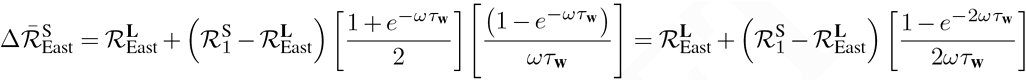

Plugging in the value of 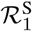:

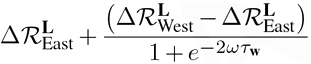

Gives:

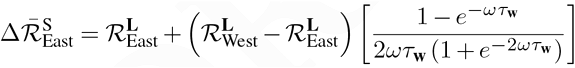

Yielding a learning time of:

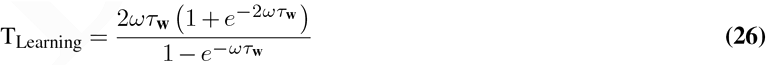

From Eq. 25, we can see that as the landmark cells become stronger, the shifts become stronger, as the animals position self-estimate becomes more heavily weighted toward whichever landmark it most recently saw. From Eq. 26 we see that, as landmark cells become stronger, the learning rate slows down, as landmark cells mostly see their own self-estimates; the contribution to position self-estimate from spatially disjoint landmarks decays quickly after the animal moves into the landmark firing field.

## Supplementary Note E: Periodicity of 2D representation

When 2D attractor dynamics yield a family of steady hexagonal bump patterns, this periodicity can be represented mathematically on the neural sheet as:

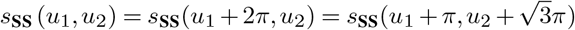

Where we have defined units on the neural sheet in terms of this periodicity. Therefore, the coordinate 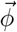 specifying a point on the manifold of stable attractor patterns is a periodic variable defined modulo the periodicity of the steady state pattern:

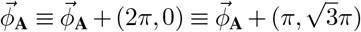

## Supplementary Note F: Proof that Learning Dynamics Reduce to a Mechanical Framework

We show how these dynamics can be reduced to a low-dimensional model where the exploration behavior gives an emergent interaction between the learned states of landmark cells, even those with non-overlapping firing patterns. We first need to solve for an animals position self-estimate as a function of its path history. To do so we first make the bookkeeping substitution:

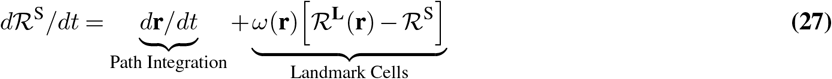

Where *ω*(**r**) = Σ*ω***H**_*i*_ (**r**) is the combined strength of all landmark cells that fire at **r**, and 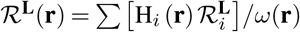 is the average position estimate being reinforced at position **r**. As the animal moves around the environment, the position self-estimate will get pushed to the learned positions of landmarks the animal visits, path integrated as the animal moves, and eventually forgotten as the animal orients itself to new landmarks. We can take this basic intuition and turn it into a closed-form equation(Verified in App. A); given any path history **r**(*t*) the solution for Eq. 27 is:

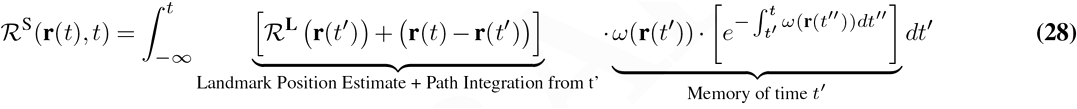

### Solving for learned position estimates as a function of current landmark position estimates

We now need to compute the mean position-self estimate seen by each landmark cell. We note that for any *individual* path, 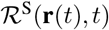 is linear with respect to 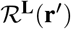. Therefore, averaging over *all* paths 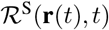 that end at **r**, the average 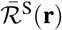 is *also* linear with 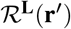. Therefore, we can construct a matrix equation:

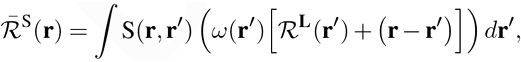

where our matrix entries S(**r**, **r′**) represent all possible ways the landmark position-estimates at position **r′** contribute to the mean position self-estimate at **r**. As long as the exploration dynamics are reversible, i.e., for any **r**(*t*), the reverse path **r**(−t) is equally likely, S is symmetric (S(**r**,**r′**) = S(**r′**,**r**)) (Proof in App. B).

To solve for the learning dynamics, we expand *ω*(**r**), 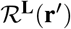 to understand the average position self-estimate as a function of position:

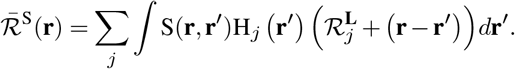

The mean position self-estimate seen by each landmark cell is then:

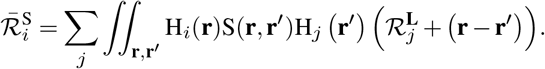

Combining this with the angle learning rule gives:

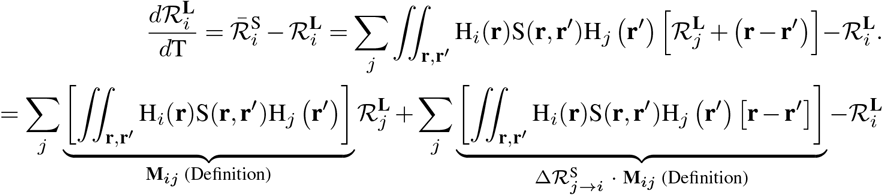

We note that Σ_*j*_ **M**_*ij*_ = 1 For all *i*^4^; therefore, we can rewrite the above equation as:

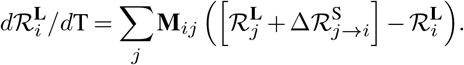

Due to the symmetry of S, **M**_*ij*_ = **M**_*ji*_, 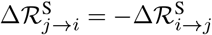. Therefore, the long term dynamics of mapping are equivalent to the first-order dynamics of a set of particles *i*, attached by springs of strength **M**_*ij*_, having a rest displacement of 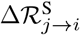. The spring constant is related to the frequency with which the animal moves between each landmark field, while the rest displacement is related to the distance between the landmark field.

### A. Proof of convolutional integral

We can check that the solution for the position self-estimate Eq. 28:

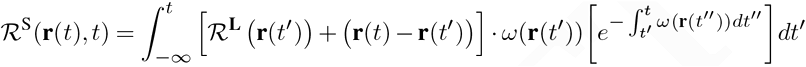

Satisfies the dynamics of Eq. 27:

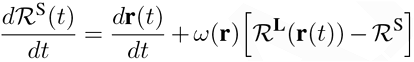

by inspection. We plug Eq. 28 into Eq. 27 to get:

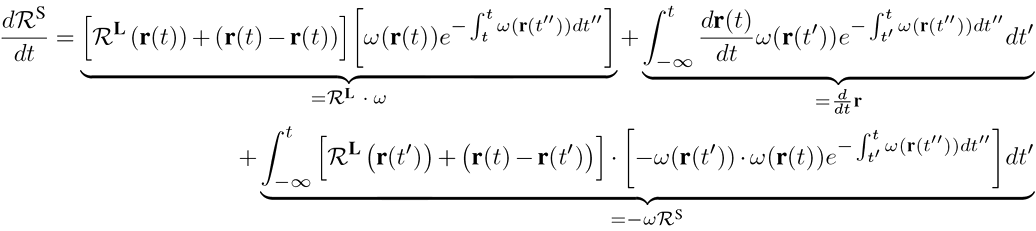

The underbraced identities are more easily seen by simplifying terms:

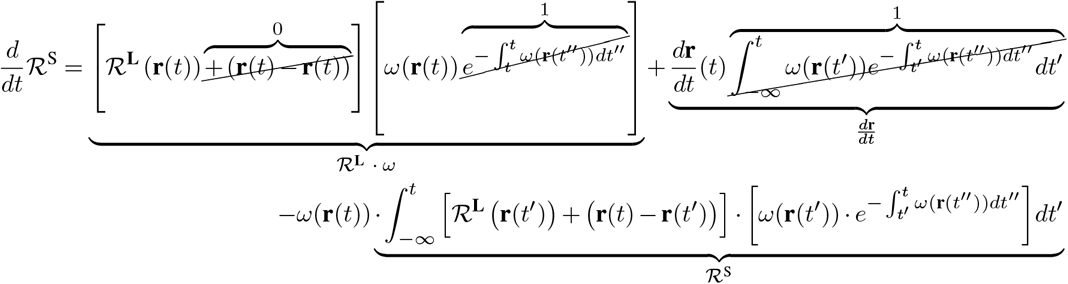

**Fig. 9.**
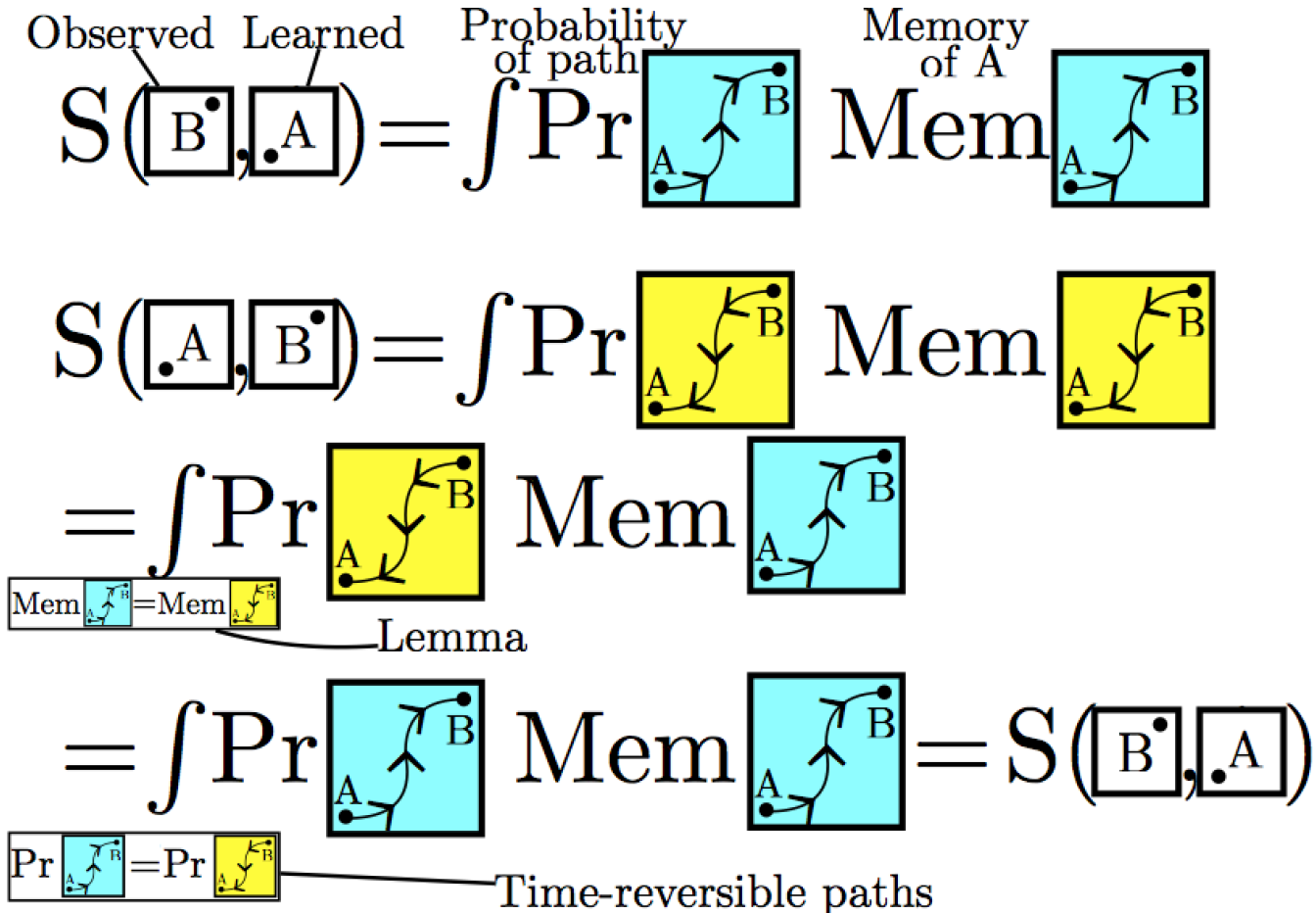
Sketch of proof in App. B that S is symmetric for time-symmetric path distributions.

### B. Proof that S is symmetric for time-symmetric path distributions

The mean position self-estimate of the animal at position **r**_B_ is the average self-estimate of all paths that pass **r**_B_ at time *t* = 0. (We pick t = 0 for mathematical convenience).

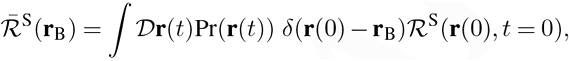

To avoid clutter, use the shorthand:

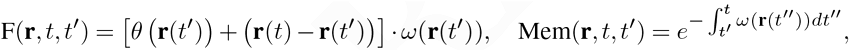

And decompose this into contributions from different past times *t′*.

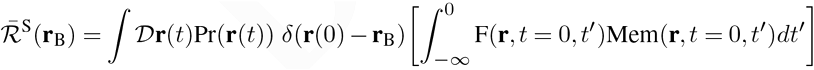

Reshuffling the order of integration and breaking things down further into contributions of **r**_A_ = **r**(*t′*)

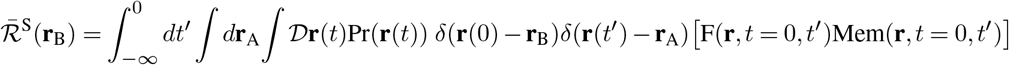

Because we have assumed the statistics of the animal trajectories r(t) will be time-reversal symmetric, the reverse, time shifted path 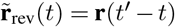 is equally likely. We therefore apply the symmetrization procedure:

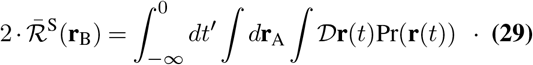

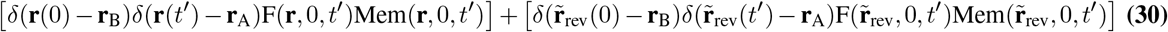

We note that Mem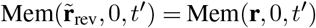, and that:

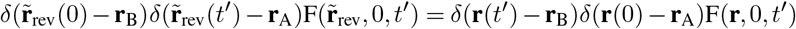

Therefore, we can simplify Eq. 30 to:

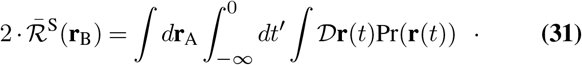

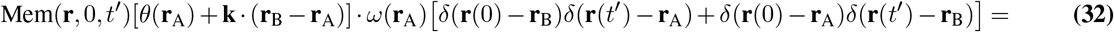

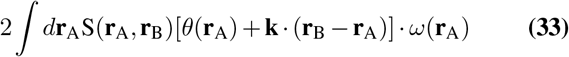

Where our matrix entries:

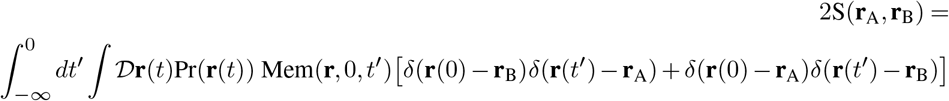

are symmetric with respect to the swapping of **r**_B_, **r**_A_.

This proof assumes uniform density of animal positions with uniform areas and strengths of each landmark cell. The proof can be generalized beyond these constraints by making effective particles corresponding to certain landmarks more “massive”, but here we present the simpler proof in the interest of clarity.

## Supplementary Note G: Simulations

### A. Exploration

In our simulations, we discretize space onto a grid. For simplicity, we have the animal follow diffusive dynamics, implemented through a random walk; at every time step, the animal moves to one of four neighboring cells; any move which would take the animal outside the box is prohibited. The animal has a position self-estimate 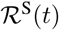 as well as an attractor state 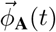, which undergoes discrete path-integration at every time step:

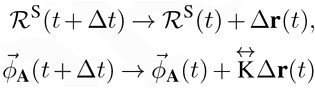

Afterwards, the position self-estimate is pulled towards the position estimates of any landmark cells which are firing:

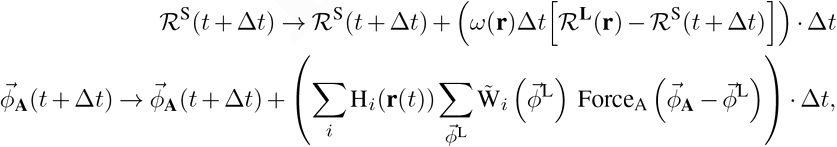

Where 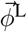 is discretized into a 15 × 15 grid so that 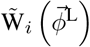 can be represented as an array.

We set the timescale of animal motion to be l

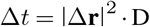

Which removes dependence on the discretization size.

### B. Learning

The learned states are initialized to their firing field center of masses. At every learning epoch **T**, the simulated animal is placed in the box with an initial position and position self-estimate and explores to get good statistics. 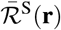, is logged, and at the end of each learning epoch, the position estimate of each landmark cell *i* is updated to be the average position self-estimate when the landmark cell is firing.

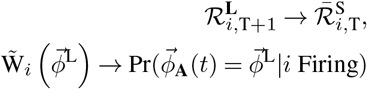

Each of these will converge after a handful of learning epochs; in practice, we use twenty.

### C. Simulation of Square and Bent Environments

Landmark cell firing fields are heterogeneous; while some are distributed across an entire border; two replicate this distribution we have two types of landmark cells in our model. 1) Landmark cells having uniform wall-length firing field, with a width of 10cm, for example 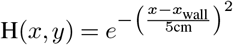 for a landmark cell on the west wall. 2) More localized, overlapping, firing fields along each wall. Each firing field is a 5 cm × 10 cm half-ellipse of along a particular wall; i.e. 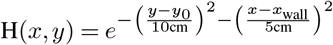 for a landmark field along the EW wall with center *y*_0_. Each type of landmark cell is evenly distributed along each wall, with the total strength and number set such that total firing strength of localized and non-localized cells is the same, and their combined strength leads to a forgetting time of *ω* = 8Hz along each wall.

Grid spacing is chosen to be 30 cm for square environments(1 ω 1 meter); We set the diffusive constant D to be (10 cm)^2^/s such that it takes an animal ~100 seconds to traverse the width of the environment.

Grid spacing is 50 *cm* with a 7° offset for the first trapezoidal environment (1.9 × .8 meters, same geometry as (11)); We set the diffusive constant D to be (20 cm)^2^/s such that it takes an animal ~100 seconds to traverse the length of the environment.

Grid spacing is 50 *cm* with a 7° offset for the second trapezoidal environment (1.9 meters long. Two straight walls with lengths of .12 meters, .6 meters, with diagonal walls starting 1 meter from the smaller straight wall (14° angle); We set the diffusive constant D to be (20 cm)^2^/s such that it takes an animal ~100 seconds to traverse the length of the environment.

The angular offset breaks the symmetry of the trapezoidal environments, yielding bending, but is not required to yield path-dependent shifts.

### D. Simulation of Topological Environments

In order to support topological defects, the environment must be filled with rich, localized landmark cues. In order for the environment to support topological defects, cues must be rich and localized, leading to uniformly distributed landmark firing fields. To model this, we have uniformly localized landmark fields, with 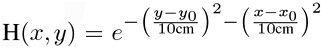, arranged at a density such that their combined strength leads to a forgetting time of 1Hz throughout the environment. The environment was 1.8 meters × 1 meter, with a center rectangular section of 1.3 × .8 meters removed. 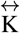 is chosen to yield a grid spacing of 60cm.

Essential ingredients to achieve topological defects are:

- A “donut-shaped” environment, which can support the topological defect.
- An environment rich in localized, strong landmark cues.
- The larger the environment is, the less deformation it has to support per unit distance, i.e. if an environment is 3 firing fields wide, a topological defect must modify the grid spacing by 33%; if the environment was 5 firing fields wide, the grid spacing would only need to be modified by 20%.
- During the “winding” procedure, the animal cannot acclimate to the intermediate environment for too long; if it fully learns the intermediate environment, the winding procedure will not work (Fig. 11).

**Fig. 10.**
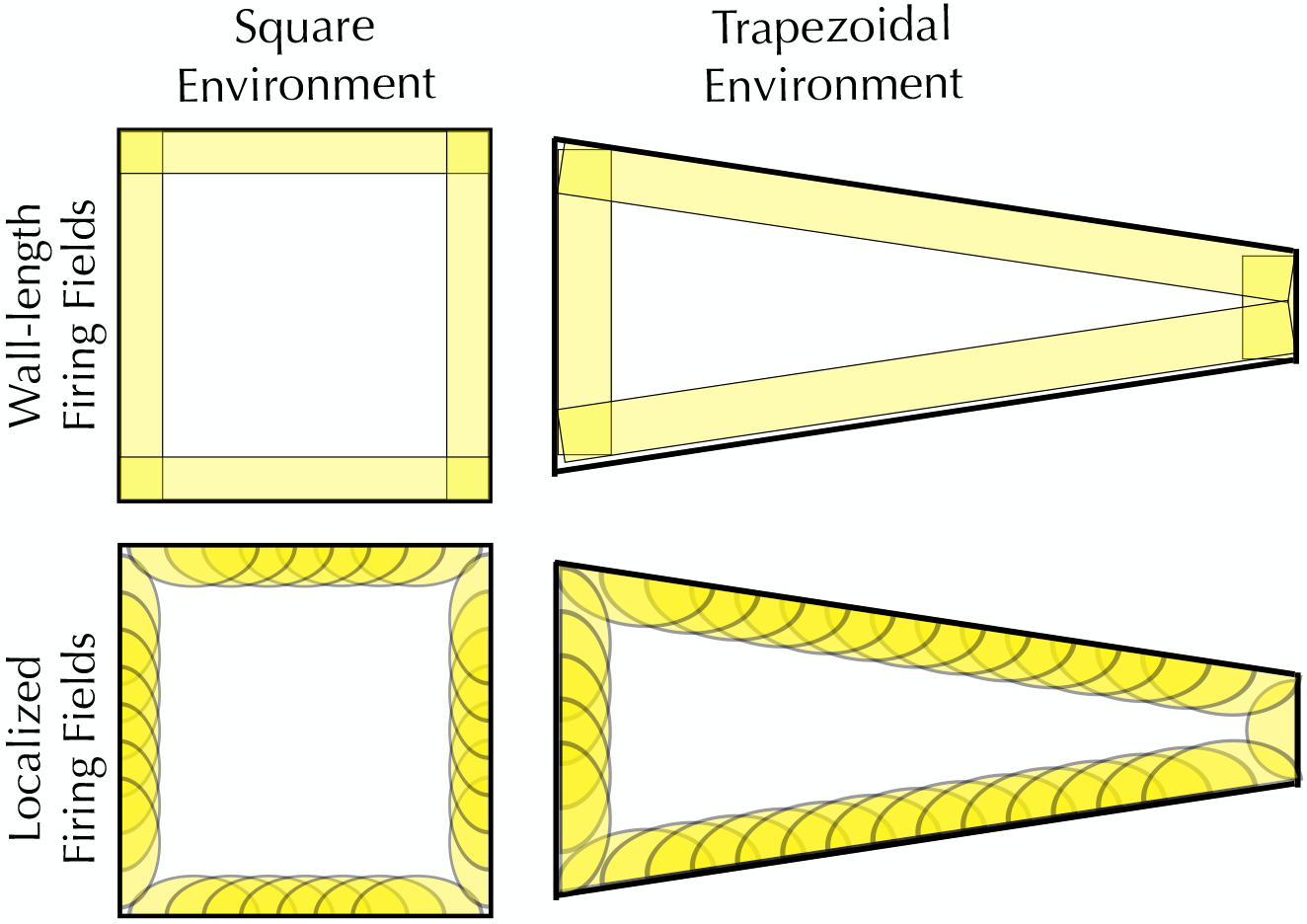
Schematic of the distribution of landmark cells for simulations of square and trapezoidal environments. To model a heterogeneous distribution of landmark cell degrees of localization, we include both landmark cells which fire uniformly along a boundary, as well as semi-elliptical landmark cells which are localized to a section of a boundary.

**Fig. 11.**
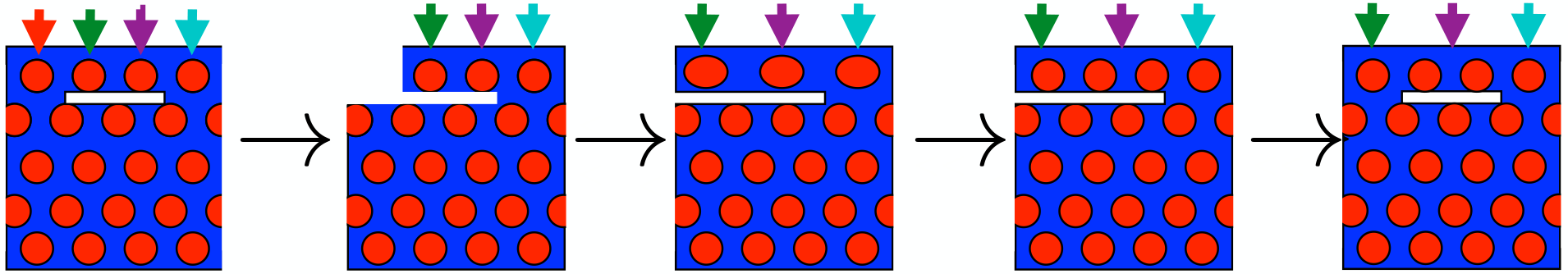
Example of topological defect failing to form dueto learning. Ifthewinding procedure is donetoo slowly, the animal will learn the deformed geometry(Third box → Fourth Box), removing the topological effect.

### E. Force law and visualization

The Grid Cell pattern is visualized by using a truncated parabolic firing rate max 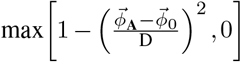, where the field width D is chosen to be 2*π*/5.

The force law chosen is a truncated sin function:

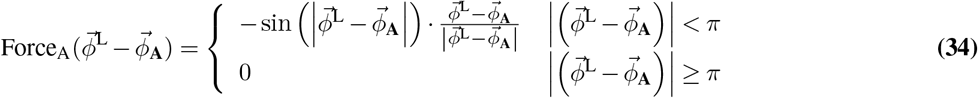

We choose this function because it has the correct qualitative features. In addition, in experimental data, the with of a firing rate peak is on the order of the spacing between two firing peaks; this prohibits a force law which is much more short-ranged than this (App. A).

**Fig. 12.**
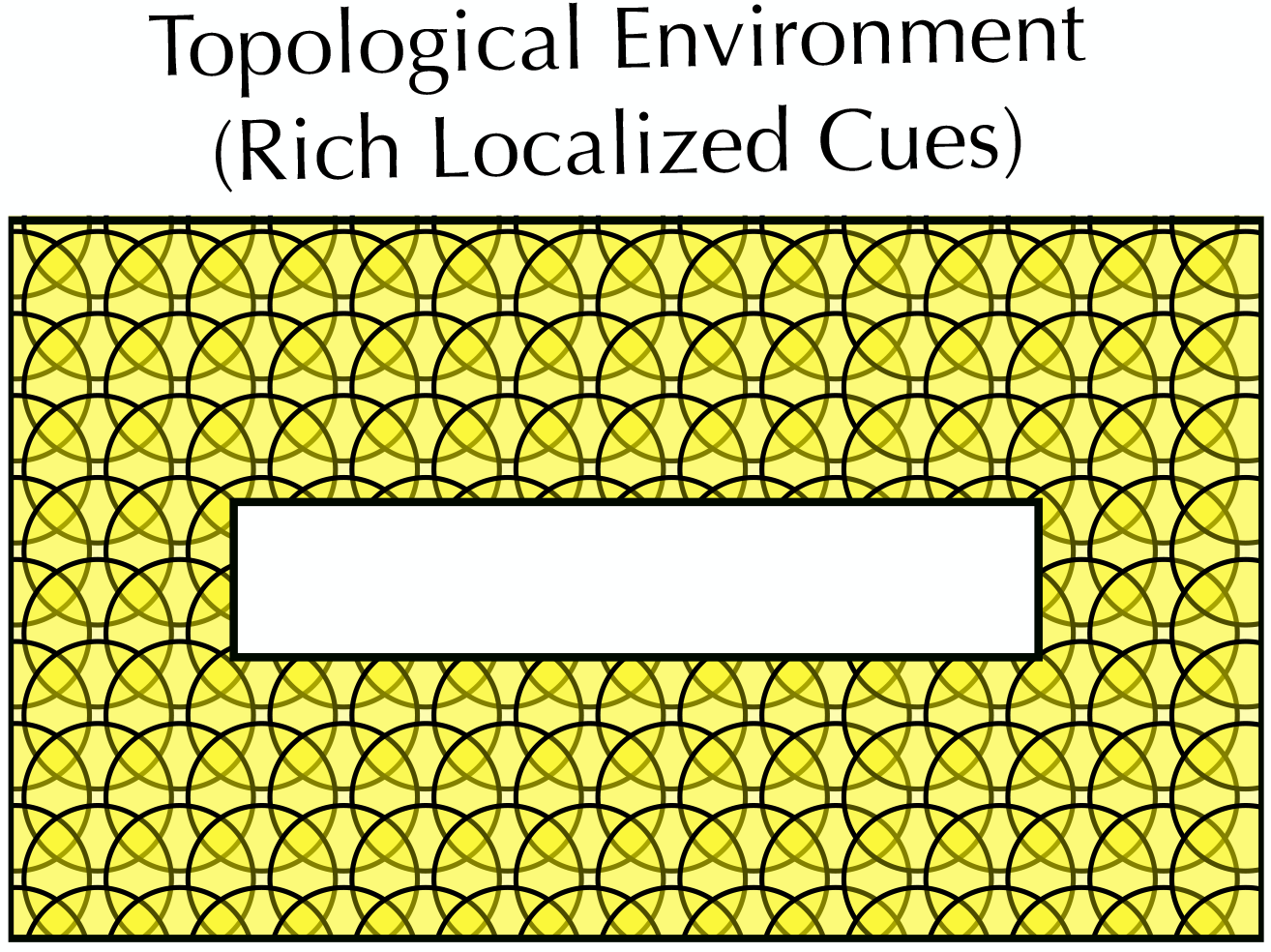
Schematic of the distribution of landmark cells for simulations the topological environment; cues are densely and uniformly localized throughout the arena.

## Supplementary Note H: Experimental Methods

Data included a subset of published neural recordings previously presented in Hardcastle et al., 2017, Hardcastle et al., 2015. Briefly, mice explored a square box while foraging for chocolate cheerios sprinkled on the floor. During each recording, neural signals from medial entorhinal cortex were recorded and subsequently clustered into distinct neurons. A grid score was computed for each cell following Langston et al., 2010. Cells above a threshold of .4 were considered grid cells. Each grid cell in the dataset was recorded after an average of 28 (data selected from Hardcastle et al., 2015) or 20 (data selected from Hardcastle et al., 2017) exposures to the recording environment.

### A. Estimation of path-dependent shifts

We examined how grid firing patterns change depending on which landmark (one of four borders in an open 1 meter box) an animal most recently encountered. To control for the effect of head direction and running speed, we preprocessed the data by translating

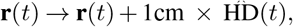

where 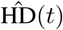 is a unit vector representing the animal’s head direction. This is to avoid artifacts related to tracking; a purely position-dependent firing rate model depends on *some* part of the animal’s body, which unlikely to be exactly the position of the tracking diode.

#### A.1. Path conditioned rate maps

We constructed maps of firing rate as a function of spatial position conditioned on the animal having last touched the North wall more recently than it touched the South Wall, etc. An animal was defined to have “touched” a wall when the head-tracking diodes came within 10cm of the wall. Varying this distance did not significantly effect our results. We avoid any sort of smoothing to prevent artifacts which might show up an experimental signature; as such, the bin size of the computed 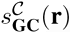 is 5cm× 5cm, and each individual trial leaves many bins for which 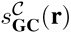 is not defined. We can create finer-grained cross-correlelograms by choosing bin sizes of 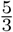 cm, and smoothing in the manner of (7), but these maps are not used for showing statistical significance. A sort of cross-correlation was taken, using the “angle” between two path-conditioned rate maps.

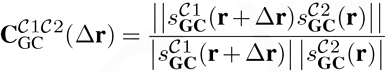

Where the mean firing rate is subtracted, and the inner product is only calculated using bins where there is data. To show significance, we calculate

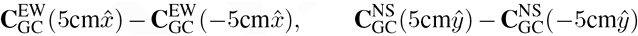

And show that the patterns are shifted towards whichever wall the animal last touched for both the EW Walls 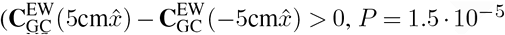, Binomial Test, *P* = 1.5 · 10^−5^, Sign-Flip Test), and the NS walls 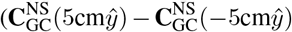, *P* = 10^−7^, Binomial Test, *P* = 10^−7^, Sign-Flip Test).

#### A.2. Spike Displacement

Starting from an adaptively smoothed firing rate map, we calculate firing field centers. For each firing field center, we gather positions of spikes recorded in that neighborhood, comparing the average spike position in that neighborhood with the firing center.

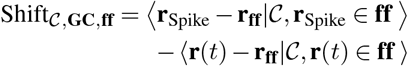

For each path condition *C*, and each firing field center **r**_**ff**_, we calculate the average spike position **r**_Spike_ within that firing field, and subtract the average *mouse* position **r**(*t*) within that firing field. The animal’s position within the firing field is subtracted to eliminate any systematic biases that might come from the animal trajectory rather than the actual neural activity (Fig. 14).

**Fig. 13.**
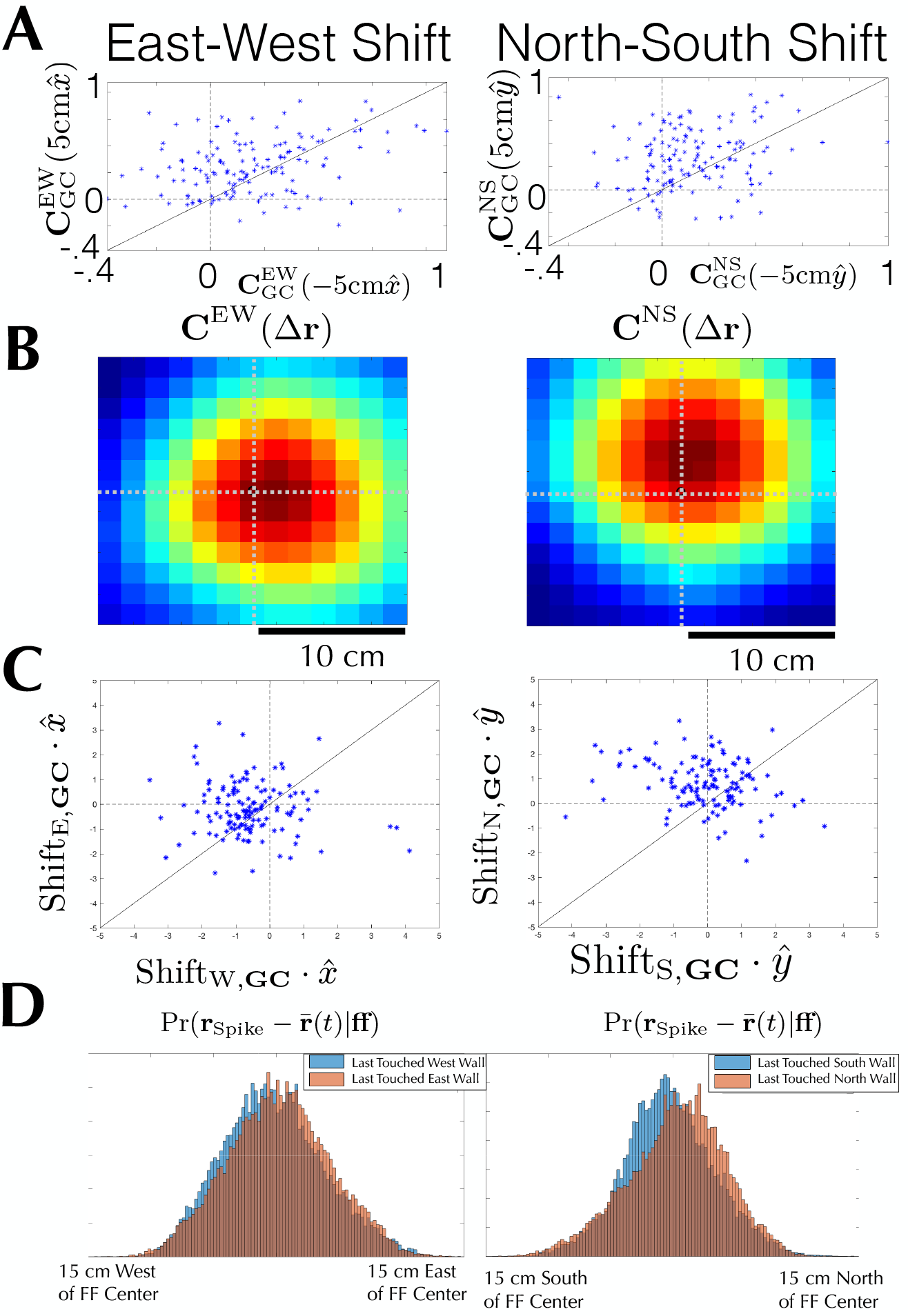
Path-dependent shifts demonstrated by Cross-Correlograms of individual grid cells. Most cells fall on the upper left of the plots, showing that the patterns tend to be shifted towards whichever wall the animal last touched for both the the EW Walls (*P* = 1.5 · 10^−5^, Binomial Test, *P* = 1.5 · 10^−5^, Sign-Flip Test), and the NS walls (*P* = 10^−7^, Binomial Test, *P* = 10^−7^, Sign-Flip Test). **B)** The path-dependent shifts is best visualized through the Cross-Correlogram averaged over all grid cells. **C)** Path-dependent shifts demonstrated by Cross-Correlograms of individual grid cells. Most cells fall on the upper left of the plots, showing that the patterns are shifted towards whichever wall the animal last touched for both the EW Walls (*P* = 3 · 10^−4^ Binomial Test, *P* = 2 · 10^−2^ Sign-Flip Test), and the NS walls (*P* = 10^−5^ Binomial Test, *P* = 10^−5^ Sign-Flip Test). **D)** The path-dependent shifts is best visualized through a histogram of individual spike displacements.

We calculate the path-dependent shift of an individual grid cell as the average shift of all firing fields in the center:

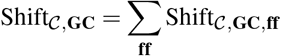

**Fig. 14.**
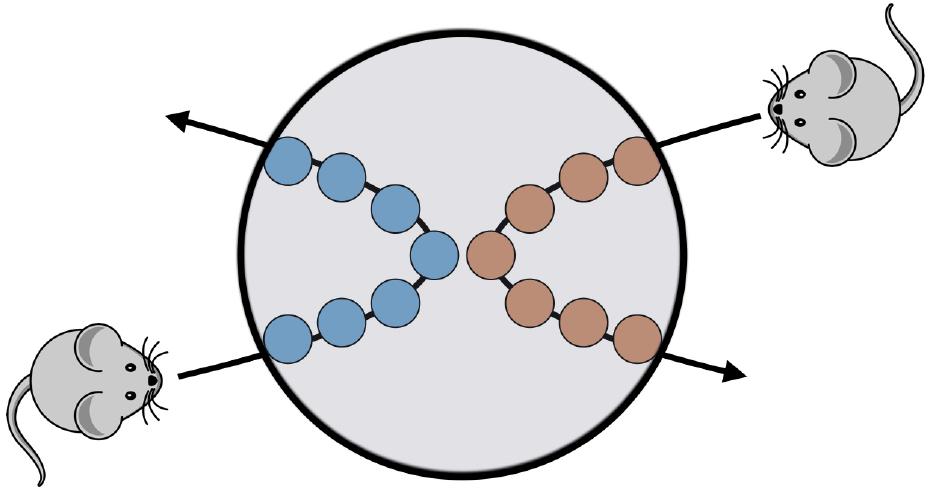
Schematic of the motivation for subtracting mouse position. An animal is most likely to be closes to the last wall it touched; if the mean animal position was not subtracted from the mean spike position, this would yield a path-dependent shift in spike positions purely dependent on animal trajectory rather than neural activity.

To show significance, for each cell, we calculate

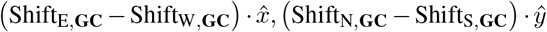

showing that the patterns are shifted towards whichever wall the animal last touched for both the EW Walls 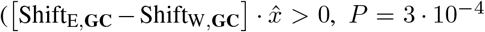, *P* = 3 · 10^−4^ Binomial Test, *P* = 2 · 10^−2^ Sign-Flip Test), and the NS walls 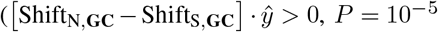, *P* = 10^−5^ Binomial Test, *P* = 10^−5^ Sign-Flip Test). We perform both binomial tests, which only depend on the sign of (Shift_E,**GC**_ − Shift_W,**GC**_), and magnitude-weighted sign-flip tests, for completeness.

## Footnotes

1 Time-reversible means that for any **r**(*t*), the reverse path **r**(−*t*) is equally likely

2 More complex, non-overlapping distributions yield the same deformations

3 We can show this by contradiction; if 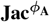 had any other non-negative modes, the family of steady states would be larger

4 This is due to the fact that shifting *all* position estimates will not change the dynamics.

## References

1. Sebastian Thrun, Wolfram Burgard, and Dieter Fox. Probabilistic robotics. MIT press, 2005.

2. Torkel Hafting, Marianne Fyhn, Sturla Molden, May-Britt B Moser, and Edvard I Moser. Microstructure of a spatial map in the entorhinal cortex. Nature, 436(7052): 801–6,2005. ISSN 0028-0836. doi: 10.1038/nature03721.

3. Edvard I. Moser, Yasser Roudi, Menno P Witter, Clifford Kentros, Tobias Bonhoeffer, and May-Britt Moser. Grid cells and cortical representation. Nature Reviews Neuroscience, 15, 06 2014.

4. Trygve Solstad, Charlotte N Boccara, Emilio Kropff, May-Britt B Moser, and Edvard I Moser. Representation of geometric borders in the entorhinal cortex. Science, 322 (5909):1865–8, 2008. ISSN 0036-8075. doi: 10.1126/science.1166466.

5. Emilio Kropff, James E Carmichael, May-Britt Moser, and Edvard I Moser. Speed cells inthe medial entorhinal cortex. Nature, 523(7561):419–424, 2015.

6. Francesca Sargolini, Marianne Fyhn, Torkel Hafting, Bruce L McNaughton, Menno P Witter, May-Britt Moser, and Edvard I Moser. Conjunctive representation of position, direction, and velocity in entorhinal cortex. Science, 312(5774):758–762, 2006.

7. Tor Stensola, Hanne Stensola, May-Britt Moser, and Edvard I. Moser. Shearing-induced asymmetry in entorhinal grid cells. Nature, 518, 02 2015.

8. Caswell Barry, Robin Hayman, Neil Burgess, and Kathryn J Jeffery. Experience-dependent rescaling of entorhinal grids. Nature neuroscience, 10(6):682–684,2007. ISSN 1097-6256. doi: 10.1038/nn1905.

9. Caswell Barry, Lin Ginzberg, O’Keefe, John, and Neil Burgess. Grid cell firing patterns signal environmental novelty by expansion. Proceedings of the…, 109(43): 17687–17692,2012. ISSN 0027-8424. doi: 10.1073/pnas.1209918109.

10. Francis Carpenter, Daniel Manson, Kate Jeffery, Neil Burgess, and Caswell Barry. Grid cells form a global representation of connected environments. Current Biology, 25(9):1176–1182, 2018/04/04 2015. doi: 10.1016/j.cub.2015.02.037.

11. Julija Krupic, Marius Bauza, Stephen Burton, Caswell Barry, and O’Keefe, John. Grid cell symmetry is zenvironmental geometry. Nature, 518(7538), 2015. ISSN 0028-0836. doi: 10.1038/nature14153.

12. Julija Krupic, Marius Bauza, Stephen Burton, and John O’Keefe. Local transformations of the hippocampal cognitive map. Science, 359(6380):1143–1146, 2018. ISSN 0036-8075. doi: 10.1126/science.aao4960.

13. Julija Krupic. Brain crystals. Science, 350(6256):47–48, 2015. ISSN 0036-8075. doi: 10.1126/science.aad3002.

14. Julija Krupic, Marius Bauza, Stephen Burton, and John O’Keefe. Framing the grid: effect of boundaries on grid cells and navigation. The Journal ofphysiology, 594(22): 6489–6499, 2016.

15. Julija Krupic, Marius Bauza, Stephen Burton, Colin Lever, and O’Keefe, John. How environment geometry affects grid cell symmetry and what we can learn from it. …of the Royal…, 369(1635):20130188, 2014. ISSN 0962-8436. doi: 10.1098/rstb.2013.0188.

16. Bruce L McNaughton, Francesco P Battaglia, Ole Jensen, Edvard I Moser, and May-Britt Moser. Path integration and the neural basis of the cognitive map. Nature Reviews Neuroscience, 7(8):663–678, 2006.

17. Talfan Evans, Andrej Bicanski, Daniel Bush, and Neil Burgess. How environment and self-motion combine in neural representations of space. The Journal of Physiology, 594(22):6535–6546, 2016. doi: 10.1113/JP270666.

18. Florian Raudies, James R Hinman, and Michael E Hasselmo. Modelling effects on grid cells of sensory input during self-motion. The Journal of physiology, 594(22): 6513–6526, 2016.

19. Marianne Fyhn, Torkel Hafting, Alessandro Treves, May-Britt B Moser, and Edvard I Moser. Hippocampal remapping and grid realignment in entorhinal cortex. Nature, 446(7132):190–4, 2007. ISSN 0028-0836. doi: 10.1038/nature05601.

20. Kiah Hardcastle, Surya Ganguli, and Lisa M Giocomo. Environmental boundaries as an error correction mechanism for grid cells. Neuron, 86(3):827–839, 2015.

21. Lisa M Giocomo. Environmental boundaries as a mechanism for correcting and anchoring spatial maps. The Journal of Physiology, 594(22):6501–6511, 11 2016. doi: 10.1113/JP270624.

22. Alex T. Keinath, Russell A. Epstein, and Vijay Balasubramanian. Environmental deformations dynamically shift the spatial metric of the brain. bioRxiv, 2017. doi: 10.1101/174367.

23. Samuel Ocko, Kiah Hardcastle, Lisa Giocomo, and Surya Ganguli. Evidence for optimal bayesian cue combination of landmarks and velocity in the entorhinal cortex. Cosyne, 2017.

24. Michael J Milford, Gordon F Wyeth, and David Prasser. Ratslam: a hippocampal model for simultaneous localization and mapping. In Robotics and Automation, 2004. Proceedings. ICRA’04. 2004 IEEE International Conference on, volume 1, pages 403–408. IEEE, 2004.

25. M Mulas, N Waniek, and J Conradt. Hebbian plasticity realigns grid cell activity with external sensory cues in continuous attractor models. Frontiers in computational…, 2016.

26. Gustavo Deco, Viktor K. Jirsa, Peter A. Robinson, Michael Breakspear, and Karl Friston. The dynamic brain: From spiking neurons to neural masses and cortical fields. PLOS Computational Biology, 4(8):1–35, 08 2008. doi: 10.1371/journal.pcbi.1000092.

27. Alexei Samsonovich and Bruce L. McNaughton. Path integration and cognitive mapping in a continuous attractor neural network model. Journal of Neuroscience, 17 (15):5900–5920, 1997. ISSN 0270-6474. doi: 10.1523/JNEUROSCI.17-15-05900.1997.

28. Yoram Burak and Ila R Fiete. Accurate path integration in continuous attractor network models of grid cells. PLoS Comput Biol, 5(2):e1000291, 2009.

29. Guifen Chen, Daniel Manson, Francesca Cacucci, and Thomas Joseph Wills. Ab-sence of visual input results in the disruption of grid cell firing in the mouse. Current Biology, 26(17):2335–2342, 2016.

30. KM Gothard, WE Skaggs, and Bruce L. Dynamics of mismatch correction in the hippocampal ensemble code for space: interaction between path integration and environmental cues. J. Neurosci., 16(24):8027–40,1996. ISSN 0270-6474.

31. Kiah Hardcastle, Niru Maheswaranathan, Surya Ganguli, and Lisa M. Giocomo. A multiplexed, heterogeneous, and adaptive code for navigation in medial entorhinal cortex. Neuron, 94(2):375 – 387.e7, 2017. ISSN 0896-6273. doi: https://doi.org/10.1016/j.neuron.2017.03.025.

